# Alpha-synuclein oligomers activate NFAT proteins modulating synaptic homeostasis and apoptosis

**DOI:** 10.1101/2023.02.21.529374

**Authors:** Ricardo Sant’Anna, Bruno K Robbs, Júlia Araújo de Freitas, Patrícia Pires dos Santos, Annekatrin König, Tiago Fleming Outeiro, Debora Foguel

## Abstract

Soluble oligomeric forms of alpha-synuclein (aSyn-O) are believed to be one of the main toxic species in Parkinson’s disease (PD) leading to degeneration. aSyn-O can induce Ca^2+^ influx, over activating downstream pathways leading to PD phenotype. Calcineurin (CN), a phosphatase regulated by Ca^2+^ levels, activates NFAT transcription factors that are involved in the regulation of neuronal plasticity, growth and survival. Here, we investigate NFAT’s role in neuronal degeneration induced by aSyn-O. aSyn-O are toxic to neurons leading to cell death, loss of neuron ramification and reduction of synaptic proteins which are reversed by CN inhibition with ciclosporin-A or VIVIT, a NFAT specific inhibitor. aSyn-O induce NFAT nuclear translocation and transactivation. We found that aSyn-O modulates the gene involved in the maintenance of synapses, synapsin 1 (*Syn 1*). Syn1 mRNA and protein and synaptic *puncta* are drastically reduced in cells treated with aSyn-O which are reversed by NFAT inhibition. For the first time a direct role of NFAT in aSyn-O-induced toxicity and *Syn1* gene regulation was demonstrated, enlarging our understanding of the pathways underpinnings synucleinopathies.

## Introduction

Parkinson’s disease (PD) is characterized by a preferential loss of dopaminergic (DA) neurons in the *substantia nigra* and accumulation of intracellular inclusions known as Lewy bodies (LB) and Lewy neurites (LN). These inclusions are primarily made up of the protein alpha-synuclein (aSyn)^1^, an abundant presynaptic and nuclear protein^2,3^, possibly playing a role in vesicular trafficking and synaptic vesicle biology^4^. In Lewy body diseases, aSyn is thought to aggregate into fibrils that are the main components of LB and LN^1,5^. The accumulation of these deposits is accompanied by inflammation, progressive neuronal dysfunction and, eventually, death of affected neuronal populations, leading to the various clinical features of the disease^6–8^. Several missense mutations in the *SNCA* gene, which encodes for aSyn, are linked to rare forms of familial PD^9–15^, suggesting aSyn may gain a toxic function that possibly contributes to disease. Although aSyn and its mutants can form amyloid fibers *in vitro*, and accumulate in LBs in the brain, aSyn fibrils have been hypothesized to be protective, by sequestering the more toxic intermediate aggregates such as oligomers^16^. This is consistent with the observation that, in certain juvenile forms of PD, extensive neurodegeneration is observed in the absence of LB formation^16–18^. In addition, it has also been shown that LB-bearing neurons may not exhibit cytotoxic phenotypes^19–21^. Furthermore, *in vitro* and *in vivo* experiments have illustrated that aSyn oligomers (aSyn-O) are toxic to neural cells, while fibrils are not^22–24^. Interestingly, some aSyn mutants, such as A30P, have a higher propensity to form oligomers than fibers^25,26^, which is advantageous for testing phenotypes associated with the accumulation of oligomeric aSyn species.

An implication of the theories of spreading of aSyn pathology^5,27,28^ is the occurrence of aSyn extracellular species. In fact, such species were shown to be of pathological relevance in various experimental paradigms^29–31^. aSyn-O can be released from cultured cells and primary neurons^32–34^, and are detectable in the soluble protein fraction of brains of patients with LB dementia^35,36^, as well as in the cerebral spinal fluid of PD patients^35^. Importantly, the occurrence of extracellular aSyn-O in other synucleinopathies suggests they may be relevant not only in PD but also in other related disorders^29^.

One of the possible effects of extracellular oligomeric forms of aSyn is the induction of Ca^2+^ influx into neurons and glial cells, and the consequent activation of downstream pathways, leading to pathological alterations and, ultimately, to cell death^37–39^.

Calcineurin (CN) is largely responsible for transducing signals generated by changes in Ca^2+^ levels^40^. CN is a calmodulin-dependent serine/threonine phosphatase. Pharmacological inhibition of calcineurin catalytic subunit (calcineurin A; CnA), with a clinically used inhibitor such as FK506 can reduce aSyn toxicity in different models^37,38^. The family of transcription factors (TF) named nuclear factor of activated T-cell (NFAT), which includes NFATc1, NFATc2, NFATc3 and NFATc4, is the main family of TF downstream of CN^41^. In resting cells, NFAT proteins are hyperphosphorylated and reside mainly in the cytoplasm. Upon activation, NFAT undergoes rapid CnA-dependent dephosphorylation and translocate to the nucleus, where they regulate gene transcription^42^. The immunosuppressive drugs cyclosporin A (CsA) and FK506, which are well-known inhibitors of CnA, inhibit NFAT activity by blocking its dephosphorylation^43,44^. More recently a new and specific NFAT inhibitor called VIVIT has been described inhibiting only the interaction of CN/NFAT, but not inhibiting the activity of the enzyme^45,46^. Although the family of NFAT transcription factors was initially characterized in the immune system, recent studies have highlighted the importance of this family of proteins in neurons, where they are involved in the regulation of synaptic plasticity, axonal growth, and neuronal survival^47^. NFAT proteins directly regulate the expression of apoptosis-related genes such as *Fas-L, Nur77, c-Flip* and with the promotion of inflammatory process such as *TNF-*α, *COX-2* and *IL1-*β in different cell types^42,48,49^. Both, apoptosis and inflammation are processes widely involved in the development of PD. However, the involvement of NFAT in aSyn-mediated degeneration of mesencephalic DA (mDA) neurons is unclear.

Here, we investigated the role of NFAT signaling in the cellular pathology and loss of synaptic proteins associated with extracellular aSyn. We demonstrate that aSyn-O induce cell death in primary DA neurons and neuroblastoma cells in a CN/NFAT dependent fashion, since it’s inhibition completely abolishes the toxic effect of the oligomers. Accordingly, after the analysis of a panel of 91 genes related to various cellular processes involved in PD/synucleinopathies by real-time qPCR, and further validation at protein level, we found that aSyn-O modulate the gene involved in the maintenance of synapses, synapsin 1 (*Syn1*). Syn1 mRNA and protein and synaptic *puncta* are drastically reduced in cells treated with aSyn-O. Furthermore, we show for the first time the presence of two conserved putative NFAT biding site in the promoter regions of *Syn1* gene and that NFAT specific inhibition rescues synapsin 1 transcripts and protein to normal levels. To our knowledge, this is the first demonstration of a direct role of NFAT in aSyn-O-induced toxicity and *Syn1* gene regulation, with implication to synucleinopathies.

## Materials and Methods

### A30P-aSyn purification

Purification of A30P-aSyn mutant was performed according to Conway *et al*., 1998^15^. The cDNA encoding the aSyn was cloned into the expression plasmid (pET21) and transformed by heat shock into the *Escherichia coli* strain (BL21DE3). Bacteria were cultured at 37 ° C under shaking at 180 rpm in the presence of 100 μg/mL ampicillin until reaching the Optical Density (OD) of 0.6 at 600 nm. Induction of the expression of the protein was performed using 500 mM isopropyl 1-thiol-β-galactopyranoside (IPTG) (Bioagency). Cells were then kept under the same conditions for 4 h and collected by centrifugation at 6,000 rpm for 15 min at 4° C. The supernatant was discarded, and the pellet resuspended in lysis buffer (50 mM Tris, 50 mM KCl). The solution was sonicated for 30 min/30 sec intervals at 4° C. Cell debris was removed by centrifugation at 13,000 rpm for 20 min at 4° C. The sample was then boiled for 15 min and centrifuged at 13,000 rpm for 20 min. After centrifugation, the volume of the supernatant was measured and transferred to a becker, added 361 g/L of (NH_4_)_2_SO_4_, stirred for 30 min and again centrifuged. In order to remove salt, the precipitate was resuspended in 25 mM Tris buffer pH 7.5 and dialyzed 4 times against H_2_O milliQ in a final volume of 4 L at 4° C on a 3,500 Da pore membrane (Spectrum™ Spectra/Por™ Fischersci). Subsequently, the sample was applied to an ion exchange column (Source 30Q GE Healthcare) initially equilibrated with 25 mM Tris buffer pH 7.5, and a 0 to 1 M NaCl gradient was performed. The presence of protein was verified by the absorbance of the samples at 280 mm. The samples corresponding to aSyn were collected, dialyzed against H_2_O milliQ, aliquoted into 1 mL tubes and lyophilized for 18 h. The samples were then resuspended in phosphate buffer (PBS) and loaded onto an 18% SDS-PAGE gel. In addition, prior to use, the samples were subjected to a new purification step by exclusion chromatography on a Superdex 200 column (GE Healthcare®). The system was equilibrated with PBS buffer at a flow rate of 0.5 mL/min. The elution was monitored by absorbance at 280 nm and the peak corresponding to the monomer was collected and used in all assays.

The samples were concentrated by ultrafiltration in an AMICON system (Millipore) with a 10,000 Da exclusion membrane. The concentration of aSyn samples was estimated by measuring the absorbance at 280 nm using the extinction coefficient of 5,960 M^-1^cm^-1^.

### A30P-aSyn oligomerization and aggregation

After purification and confirmation of the purity of the sample, aSyn was incubated at the concentration of 140 μM in PBS pH 7.5 under sterile conditions, under agitation at 800 rpm and 37° C in a Thermo-Shaker for TS-100 microtubes (Biosan) for up to 10 days. A 5 μL aliquot of the aSyn protein not incubated or incubated for different periods under aggregating conditions was transferred to a nitrocellulose membrane (Bio-rad®; 0.45 μm). The sample was then blocked with OdysseyTM blocking buffer for 60 min at 22° C. After this period, the membrane was incubated with primary antibody against aSyn (Monoclonal anti-human mouse α-synuclein Syn211 Invitrogen™), against A11 oligomers (Polyclonal Rabbit Anti-amyloid oligomer A11 AB9234 Millipore™) and against amyloid fibers OC (Polyclonal Rabbit Anti-amyloid fibers OC AB2286 Millipore™) all at 1:1000 dilution for 18 h at 4° C. After the incubation period, the membrane was washed 5 times under agitation for 5 min each wash with 0.1% PBS-Tween20. After washing, the membrane was incubated with antibody against IgG at a dilution of 1:10,000, in Odyssey™ blocking buffer + 0.1% PBS, which can be anti-rabbit or mouse depending on the host of the primary antibody. Incubation took place for 2 h at 22° C under gentle agitation and then the membrane was washed 5 times with 0.1% PBS-Tween20 under agitation for 5 min each wash and kept in PBS buffer protected from light. The membrane was then digitized and scanned in the appropriate channel (700 nm for Cy5.5 antibody and 800 nm for IRDyeTM antibody using the Odyssey Infrared Imaging System instrument.

### Atomic force microscopy (AFM)

Topographic AFM images were obtained at room temperature using TappingMode® AFM with Dimension ICON (Novascan). Images were acquired using Si3N4 oxide AFM (K∼3 N/m) cantilever (Model: OTESPA). The scan rate was 0.5 Hz, and the scan resolution was 512 per line. The images were subjected to baseline adjustment with 2nd order polynomial when necessary to reduce the effects of curvature and skew of the images. To obtain the images, 5 μL of samples aggregated for different periods were applied on a previously cleaved mica. The samples were in contact with the mica for 5 min and washed with 100 μL of MilliQ water. The samples adsorbed onto the mica were then incubated at room temperature for 24 h to dry.

### Detection of aSyn fibrillization using Thioflavin-T

A Thioflavin-T (Th-T) assay was performed to characterize the species of aSyn present at different times of aggregation. Th-T is a dye widely used to detect amyloid fibrils, since its fluorescence increases upon biding specifically to those structures. Fluorescence spectra of Th-T were obtained using the ISS-PC1 spectrofluorometer (ISS, Champaign, IL) with excitation at 450 nm and the emitted light was detected in the range between 470-530 nm. An aliquot of aSyn sample was mixed with Th-T in PBS buffer before measurement. The ratio between the concentration of aSyn and Th-T (Sigma) was 1:10 in all assays.

### Primary culture of dopaminergic neurons

C57BL6 animals were used from the 15th embryonic day (E15). Mice were bred and maintained under specific pathogen free conditions in the animal facility of the University Medical Center G Cottingen (G Cottingen, Germany). Animal procedures were performed in accordance with the European Community (Directive 2010/63/EU), institutional and national guidelines (license number 19.3213). Females were euthanized by cervical dislocation and pups removed and transferred to a petri dish containing PBS buffer at 22°C. With the aid of a magnifying glass, the brains were removed from the cranial vault and the region corresponding to the ventral integumentary area (VTA) was dissected to obtain dopaminergic neurons. The tissue was then dissociated, first mechanically using forceps and then with 0.03% trypsin/PBS for 5 min at 37° C. Cells were seeded in 24-well plates with polylysine coated coverslips at a density of 60,000 cells per well. Cultures were maintained in Neurobasal medium (Gibco®) supplemented with 1% B27 (Invivocell®). The experiments were performed after 14 days of neuron maturation *in vitro*.

### Neuroblastoma 2-a (N2a) cell culture

N2a cells (ATCC® CCL-131TM) originated from murine neuroblastomas were maintained in DMEM culture medium “Dulbecco’s modified Eagles medium” (Gibco ®) supplemented with 10% Fetal Bovine Serum (FBS) (Invivocell®), 4 mM of L-glutamine (Sigma®), 100 units/mL penicillin and 100 μg/mL streptomycin at 37° C in 5% CO_2_. The cells were kept under these conditions until they reached 80% confluence, where they were either transferred to a new bottle containing new medium or seeded in plates for the assays.

### Differentiation of N2a cells in dopaminergic neurons

Differentiation of N2a cells into dopaminergic neurons was performed as described by Tremblay *et al*., 2010^50^ and Latge *et al*., 2015^51^.Cells were seeded in 6-well plates for Western blot and qPCR analysis at a density of 2.0 × 10^5^ and in 24-well plates for toxicity and NFAT transactivation assays or on coverslips at a density of 1.5 × 10^3^ for immunofluorescence analysis. They were maintained for 24 h in DMEM medium supplemented with 10% FBS (Invivocell®), 4 mM L-glutamine (Sigma®), 100 units/mL of penicillin and 100 μg/mL of streptomycin at 37 °C in 5% CO_2_. After the rest period, the maintenance medium was removed and replaced by the differentiation medium, which consists of DMEM supplemented with 0.5% FBS (Invivocell®), 4 mM L-glutamine (Sigma®), 100 units/mL of penicillin, 100 μg/mL streptomycin and 1 mM N6,2′-O-Dibutyryladenosine 3′,5′-cyclic sodium monophosphate salt (dbcAMP) (Sigma®). Cells were maintained with the differentiation medium at 37 °C in 5% CO_2_ for 3 days.

### Immunofluorescence analysis

N2a cells or primary dopaminergic neurons were fixed with 4% paraformaldehyde for 20 min at room temperature, permeabilized with Triton X-100 (1% in PBS) for 4 min at room temperature and incubated with OdysseyTM blocking buffer for 1 h at the same temperature condition. N2a cells were then incubated with polyclonal anti-TH (tyrosine hydroxylase; 1:100, Millipore) and anti-MAP2 (1:100; Millipore) antibodies to probe differentiation and with anti-NFAT (NFATc3, Santa Cruz Biotechnology) for the NFAT translocation experiments in OdysseyTM blocking buffer + 0.1% Triton X-100. Primary cells from dopaminergic neurons were incubated with anti-MAP2 (1:100, Millipore) and anti-synapsin 1 (1:200; Millipore) polyclonal antibody under the same conditions. Then, they were washed with PBS and incubated with secondary antibodies conjugated to Alexa Fluor 488 (1:1,000 anti-rabbit; Molecular Probes) and Alexa Fluor 546 (1:500 anti-mouse; Molecular Probes) for 18 h at 4 °C. Cells were then washed with PBS, stained with DAPI (300 nM), washed three more times with PBS and mounted. N2a cells were photographed using an Evos ® microscope (AMG, FL Cell Imaging System), with a 20× objective, or a Fluoview FV10i (Olympus Co.) laser scanning confocal microscope, with a 100x objective. Dopaminergic neurons were imaged using the Zeiss Axioplan Microscope (LSM800, Zeiss, USA), with a 100x objective. All images of both cell types were processed using ImageJ software.

### Cell viability assays

To determine the toxicity in N2a cell lines and in primary cultures of dopaminergic neurons, a commercial kit for membrane integrity determination (based on LDH release), cytoTox-ONE™ Homogeneous Membrane Integrity Assay (Promega®) was used. The tests were performed according to the manufacturer’s instructions. After 14 days of maturation of the primary neuron cultures *in vitro*, the cells were treated with vehicle (PBS) or the stated concentrations of monomeric, oligomeric and fibrillar aSyn. When indicated, cells were pretreated for 20 min with cyclosporine (CsA) or VIVIT (Calbiochem). After 24 h of incubation at 37°C with 5% CO_2_, the culture supernatant was collected, and the protocol suggested by the manufacturer was performed. In cultures of the N2a cell line, cells were seeded in 24-well plates with coverslips at a density of 2.0 × 10^3^ cells per well. After 24 h of incubation at 37°C and 5% CO_2_, the cells were subjected to the cAMP differentiation protocol. After the differentiation period, the cells were treated as stated before for primary neurons in culture.

### Violet crystal cell proliferation assay

N2a cells were differentiated as described above in 96-well plates, 2.0 × 10^3^ cells per well, in triplicate. Then, they were treated with 10 μM of aSyn oligomers with or without previous treatment with 10 μM of VIVIT. After 24 h, the level of toxicity was assessed by crystal violet staining. Crystal violet staining consists of fixing the cells with absolute ethanol for 10 min, staining the fixed cells with 0.005% crystal violet in 20% ethanol for 10 min, washing with distilled water, solubilizing the crystal violet with methanol for 5 min and read the plate absorbance in a spectrophotometer at 595 nm.

### Analysis of cell death by propidium iodide labeling

For cell death analysis, differentiated N2a cells were grown in 6-well plates at a density of 2.0 × 10^5^ per well. Then, they were treated with 10 μM of aSyn oligomers with or without previous treatment with 10 μM of VIVIT. After 24 h of treatment, cells were trypsinized and washed once with PBS buffer. Cells were labeled with propidium iodide (PI) (5 μg/mL). Cell death analysis was determined by collecting 10,000 events by flow cytometry (FACScalibur).

### Analysis of gene expression by real-time PCR

Differentiated N2a cells were treated with 10 μM of aSyn oligomers with or without pretreatment with 10 μM of VIVIT for 16 h. CA-NFATc2-ER N2a cells (see below) were treated with vehicle or 150 nM Tamoxifen. RNA from these cultures was extracted and purified using the RNeasy Kit (QIAGEN). Quantification of purified RNA was measured using a Nanodrop-UV-visible spectrophotometer (Thermo Scientific) and RNA integrity was determined by analyzing 28S and 18S ribosomal mRNA bands on an agarose gel. The cDNA was synthesized using the KIT Quantitec Transcription KIT (QIAGEN) as recommended by the manufacturer. Real-time polymerase chain reaction (RT-PCR) was performed using SYBR Green PCR Master mix (Applied Technologies) on QuantStudio3 Real-Time PCR System (Applied Biosystems) according to the manufacturer’s instructions. The quality of the primes was tested through the analysis of the dissociation curve (melting curve). In the first Real-Time round, we used a panel of 91 genes known to be involved in the process of programmed cell death, dynamics and formation of dendritic spines and neuronal synapses with possible implications for PD. Primers were synthesized by the company Real Time Primers. The list of these genes and their sequences are presented in Supplementary material.

### NFAT nuclear translocation

N2a cells were differentiated and treated with oligomers of aSyn. After treatment, the cells were fixed with 4% paraformaldehyde and followed by the immunocytochemistry described protocol, using DAPI for nuclear labeling and anti-NFAT (NFATc3, Santa Cruz Biotechnology). Cover slides were mounted and observed on a Fluoview FV10i (Olympus Co.) laser scanning confocal microscope to reveal cellular disposition of NFAT. The quantification of the translocation was performed with ImageJ. Nuclear region was determined by the DAPI staining. The intensity of red (NFAT) signal on nucleus or on the cytoplasm was used to calculate the ratio of translocation.

### NFAT transactivation

To study the NFAT transactivation induced by aSyn oligomers, we used co-transfection experiments with the reporter plasmid pGL4.30 (Promega) that contains upstream to the Firefly luciferase sequence 3x a binding sequence for NFAT, and the plasmid pRL-SV40 (Renilla luciferase) that is under constitutive expression promoters and allows the normalization of transcription data. In these assays, the measurement of the luminescence generated by the luciferases allows comparing the degree of NFAT activation under the conditions tested. N2a cells were seeded at 70% confluence on 24-well plates and differentiation was induced. After 72 h, cells were transfected with Fugene HD (Promega) using DNA:reagent ratio 1:3. Twenty-four h later, the cells were treated with ionomycin, aSyn oligomers or pre-incubated also with CsA and VIVIT. After more 18 h, the luminescence detection assay was performed following all the manufacturer’s recommendations (Dual-Luciferase Reporter Assay Kit System E1910 (Promega).

### CA-NFATc2-ER N2a cells construction

To confirm that cell death in differentiated N2a cultures treated with aSyn oligomers is caused by activation of NFAT and to analyze gene transcription under NFAT regulation in our models, we used a N2a strain stably transduced with CA-NFATc2-ER. Briefly, retroviral plasmid pLIRES-EGFP-CA-NFATc2-ER was constructed as described in Robbs, 2013^48^. The BD EcoPack2 ecotropic packaging cell line (BD-Biosciences) was transiently transfected with a retroviral vector via calcium phosphate precipitation for 24 h. Cell-free virus-containing supernatant was collected 48 h after transfection, mixed 1:1 with fresh medium, supplemented with 8 μg/ml polybrene (FLUKA Chemie, Buchs, Switzerland), and immediately used for spin-infection (2 × 45 min at 420 g, room temperature) of 2.5 × 10^4^ N2a cells. Infected cells were incubated at 37°C for an additional 24 h. Afterwards, transduced cells were selected with G418 (Invitrogen) at 1,000 μg/ml for 14 days where all remaining cells were GFP positive. In this chimera, the estrogen receptor truncated in the N-terminal region, and responsive only to the drug tamoxifen, is linked to the C-terminal portion of NFATc2. When tamoxifen (4-hydroxytamoxifen from Invitrogen) binds to the estrogen receptor, NFATc2 is activated^48^. We treated differentiated CA-NFATc2-ER N2a cultures with different concentrations of tamoxifen.

## Statistical analysis

All statistical analysis were performed with GraphPad Prism 5.0 software (GraphPad, San Diego, CA) from at least three independent experiments. Significance was calculated using One-Way ANOVA test with Dunnett’s posttest with multiple comparison against untreated or control sample where p < 0.05 were significant. Data are presented as means ± SD.

## Results

### A30P-aSyn oligomer-induced cytotoxicity and synapses loss is mediated by the calcineurin pathway

Previously, we have demonstrated that ectopic applied recombinant aSyn aggregates are toxic to primary cultures of mDA neurons ^52,53^. Here, we used the same approach to ask if differently aged aggregates of A30P-aSyn added to neurons in culture, in the absence or in the presence of the Calcineurin A (CnA) inhibitor, cyclosporin A (CsA), would remain toxic. First, we probed which species of A30P-aSyn, namely monomers, oligomers, or fibrils, would be cytotoxic to primary dopaminergic neurons in culture. Characterization of all these specie is shown in **Supplementary Figure 1**. Consistently with previous studies, only oligomers (A30P-aSyn-O) were toxic to these cells, causing ∼45% of cell death. However, when cells were pre-incubated with CsA, there was a significant reduction in oligomer-induced cytotoxicity (**Figure 1A**). Furthermore, we quantified the number of neurons per field in those samples and, again, we found in the A30P-aSyn-O sample a reduction of 60% in the number of neurons. CsA reverted the cytotoxicity induced by A30P-aSyn-O, reducing cell death, and restoring neuronal arborization (**Figure 1B and C**).

**Figure 1.**
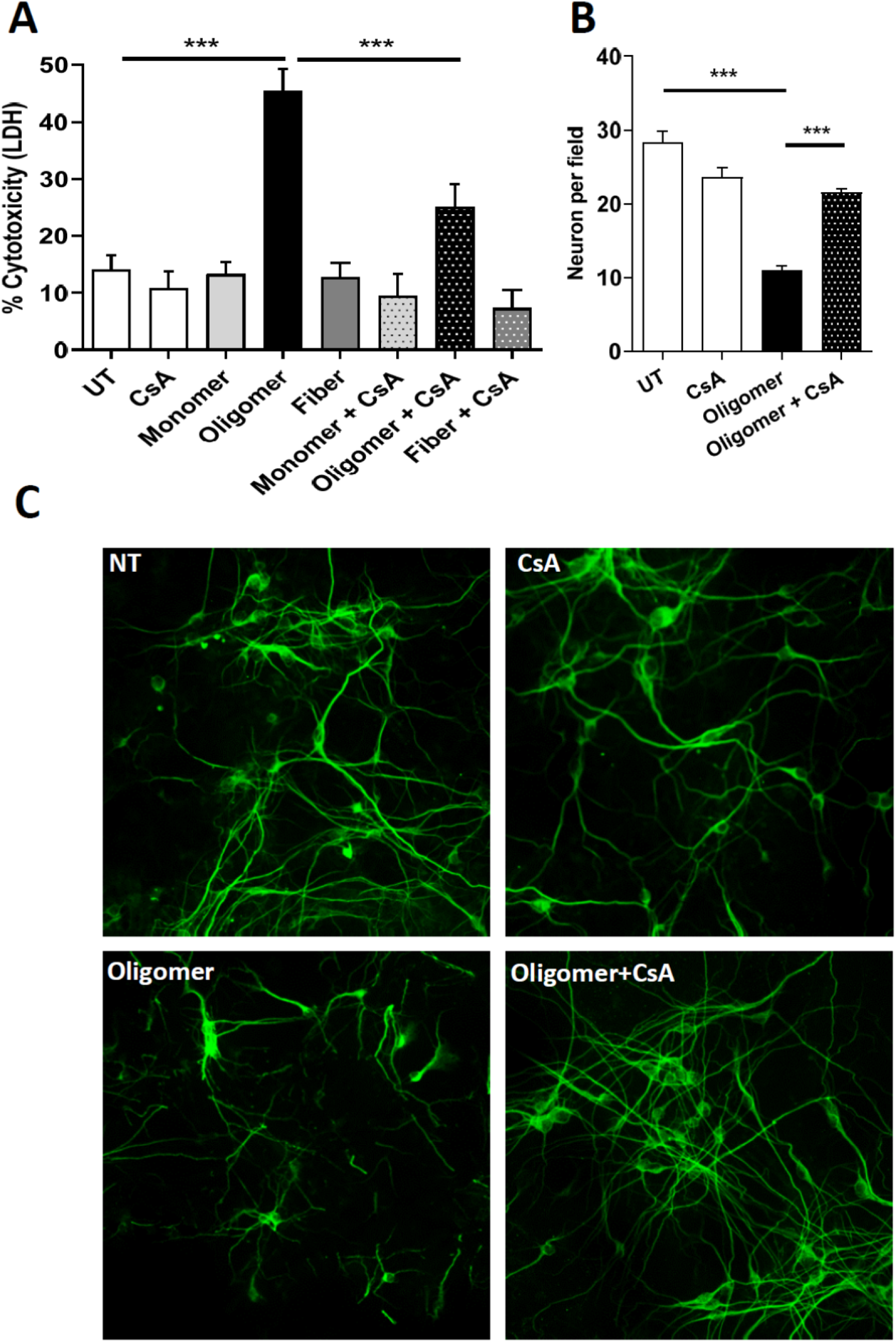
The cytotoxic effect of A30P-aSyn-O to primary dopaminergic neurons in culture is mediated by calcineurin. Mouse mesencephalic neurons were treated with 5 μM of different A30P-aSyn species (monomer, oligomer of fiber) in the absence or in the presence of 2 μM of CsA. After 24 h of treatment, cell viability was evaluated by LDH release (**A**). The number of neurons per field was quantified (**B**) in the cultures treated with A30P-aSyn-O in the presence or absence of CsA, and immuno-stained with anti-MAP2 antibody (**C**). All experiments were performed at least three times and graphs are represented as mean and SD where * p > 0.05, ** p > 0.01 and *** p > 0.001.

Next, the effects of the different species composed of A30P-aSyn on synapse formation, stabilization and morphology were investigated. It has been previously demonstrated that oligomeric aSyn reduce the levels of synapsin-1 as well as its localization in presynaptic vesicles, which correlates with cognitive impairment in patients^54^. Treatment of dopaminergic neurons with A30P-aSyn-O and fibrils, but not with monomers, led to a drastic reduction in the number of synapses containing synapsin-1 loaded vesicles. Again, this effect was completely abolished by CsA treatment (**Figure 2**). Together, these data support a major role for the calcineurin pathway in A30P-aSyn-O induced neural degeneration.

**Figure 2.**
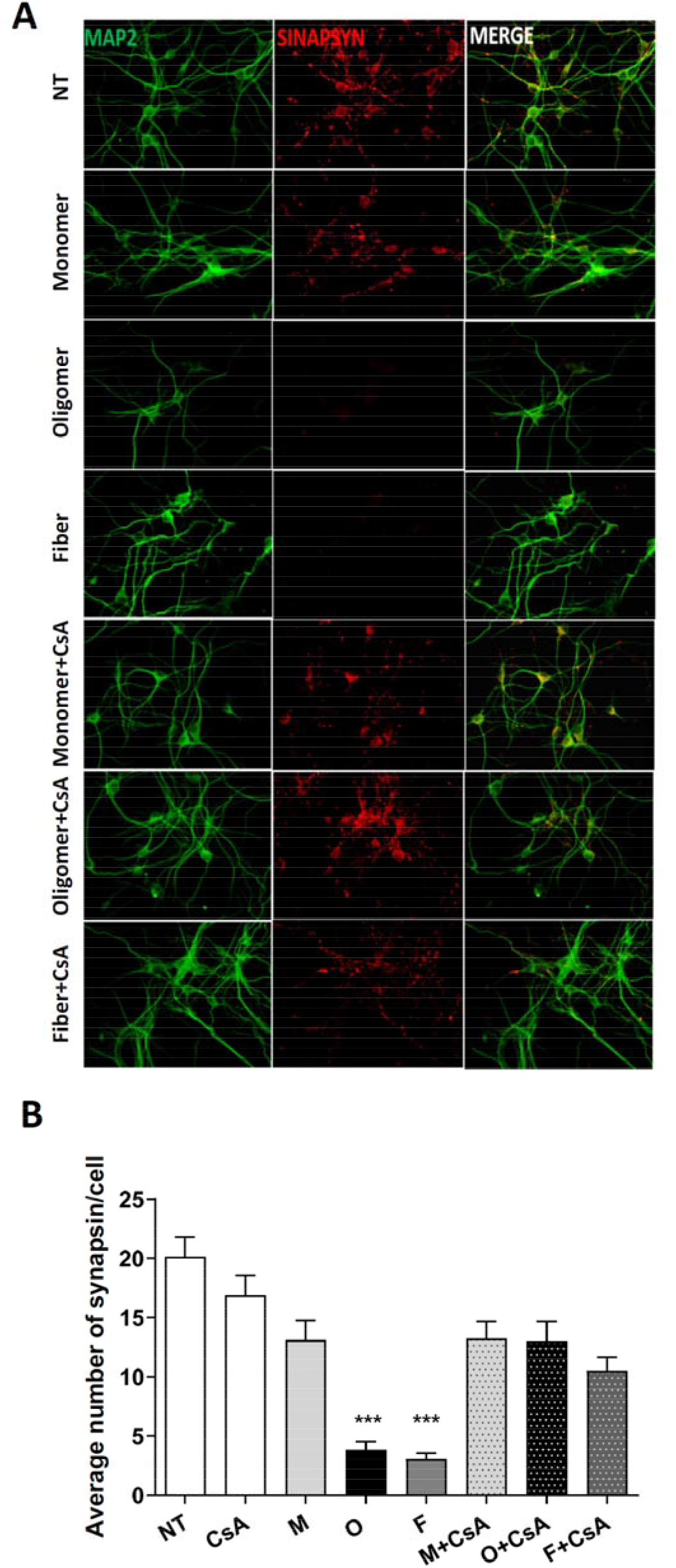
A30P-aSyn-O-induced synaptic loss in primary dopaminergic neurons is mediated by calcineurin. Primary mouse midbrain dopaminergic neurons were treated with 5 μM of aSyn monomers, oligomers or fibrils in the absence or in the presence of 2 μM of CsA for 24 h. Cells were immuno-stained with anti-MAP2 and anti-synapsin 1 antibodies (**A**). The number of synapses in each condition was evaluated by counting synaptic puncta (**B**). All experiments were performed at least three times and graphs are represented as mean and SD.

### The toxic effects of aSyn oligomers are mediated by the calcineurin-NFAT pathway leading to NFAT translocation into the nucleus activating reporter genes

To further investigate the role of CnA and the possible involvement of the NFAT transcription factor pathway in these cellular models of PD, we differentiated N2a cells into dopaminergic phenotype by treating the cells with butyryl AMPc in reduced serum medium for 72 h, as previously described^50,55^. The levels of tyrosine hydroxylase (TH) and MAP2 were evaluated as an indicator of differentiation efficacy as described in Latge *et al*., 2015^51^.

Next, we evaluated the cytotoxicity of different concentrations of A30P-aSyn monomers, oligomers, and fibrils, as previously done with primary neurons. As expected, monomers and fibrils were not significantly toxic to differentiated N2a, not even at the concentration of 10 μM, while 5 and 10 μM of oligomers were able to kill, respectively, 20% and 42% of the cells (**Figure 3A)**. Interestingly, non-differentiated N2a were insensitive to oligomers treatment at any concentration **(data not shown)** confirming the high sensitivity of dopaminergic neurons to oligomer of aSyn. Again, the pre-treatment of the culture with 2 μM of CsA significantly protected the cells from oligomer-induced toxicity **(Figure 3A)**.

**Figure 3.**
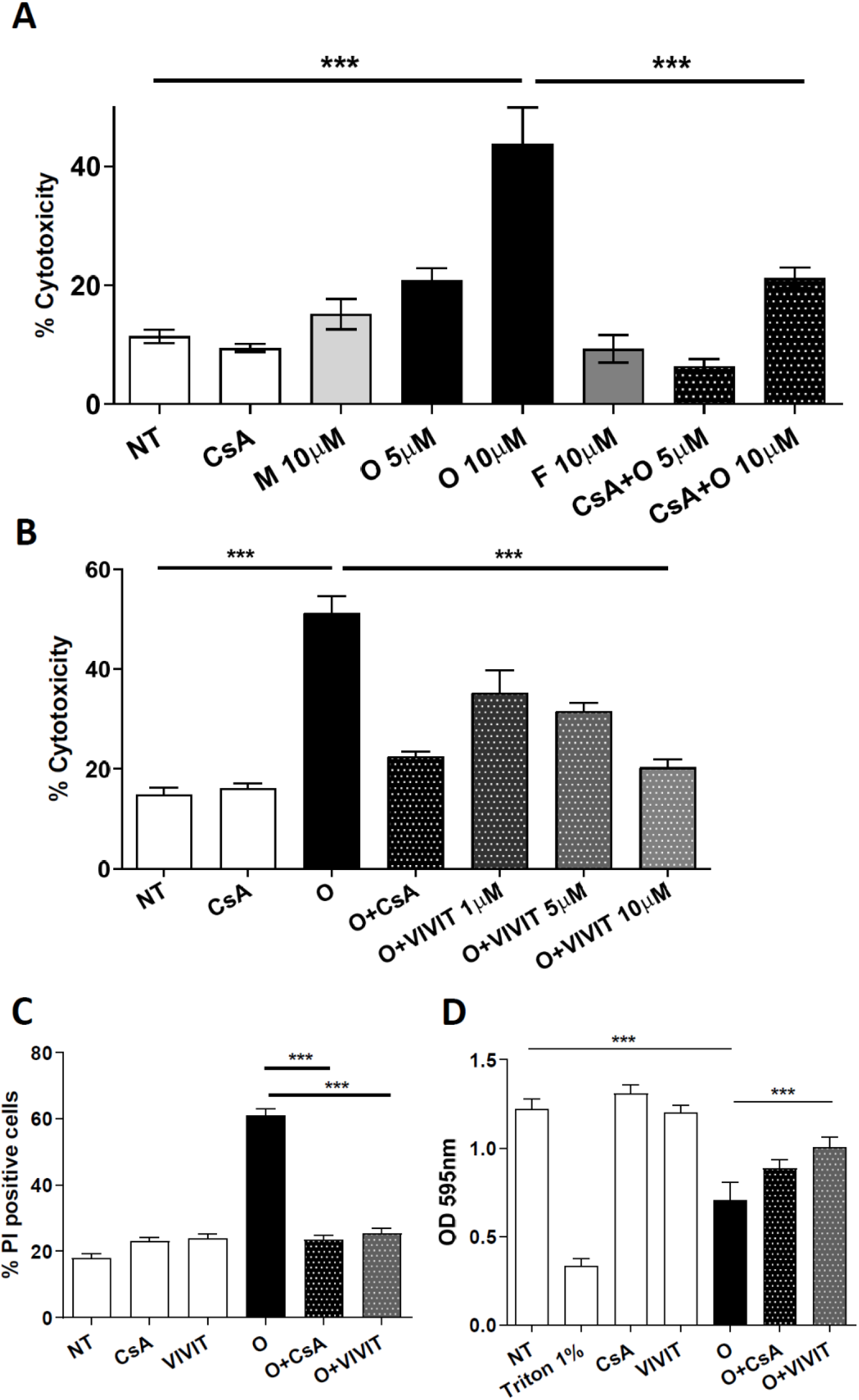
A30P-aSyn-O-induced toxicity in differentiated N2a dopaminergic cells is dependent on NFAT activation. Differentiated N2a were treated with different concentrations of A30P-aSyn monomers (M), oligomers (O), and fibrils (F) in the absence or in the presence of 2 μM of CsA and cellular damage was evaluated by LDH release (**A**). The specific NFAT inhibitor VIVIT was able to prevent the cellular damaged caused by A30P-aSyn-O evaluated by LDH (**B**), crystal violet (**C**) or propidium iodide staining (**D**). All experiments were performed at least three times and graphs are represented as mean and SD where * p > 0.05, ** p > 0.01 and *** p > 0.001.

To evaluate a direct implication of NFAT in aSyn toxicity, we used VIVIT, a selective peptide inhibitor of NFAT activation. VIVIT blocks only the interaction of CnA with NFAT, making it a very specific inhibitor of NFAT pathway, as it does not interfere with CnA phosphatase activity, as CsA^46,56^. The cells were pre-treated with 1, 5 or 10 μM of VIVIT followed by the addition of 10 μM A30P-aSyn-O, and cell death was evaluated as described **(Figure 3B)**. We observed a dose dependent inhibition of oligomer-induced cytotoxicity by the analysis of cell membrane integrity loss by propidium iodide (PI) (**Figure 3C**) and by crystal violet staining **(Figure 3D)**. Around 60 % of the cells treated with oligomers were positive for PI, while pre-treatment with VIVIT lowered PI signal to around 30%. As expected, 2 μM of CsA prevented oligomer-induced cytotoxicity. Together these data strongly support the hypothesis that the NFAT pathway is involved in neuronal cell death induced by aSyn oligomers.

To determine whether the NFAT transcription factor is activated by A30P-aSyn-O, we assessed the subcellular localization of NFAT upon treatment of the cells with oligomers. As expected, in untreated, control cells, NFAT was found mainly in the cytosol, and in almost all cells the nuclei were not stained. When cells were treated with A30P-aSyn-O, NFAT could be detected in the nucleus 3 h after treatment (**Figure 4A**). Importantly, NFAT-nuclear localization was completely abolished by treatment with CsA or VIVIT (**Figure 4A**). The extent of NFAT translocation into the nucleus was quantified by measuring the ratio between the intensities of the signal in the nucleus and in the cytosol, and we found a 3.5-fold increase upon treatment with A30P-aSyn-O **(Figure 4B)**.

**Figure 4.**
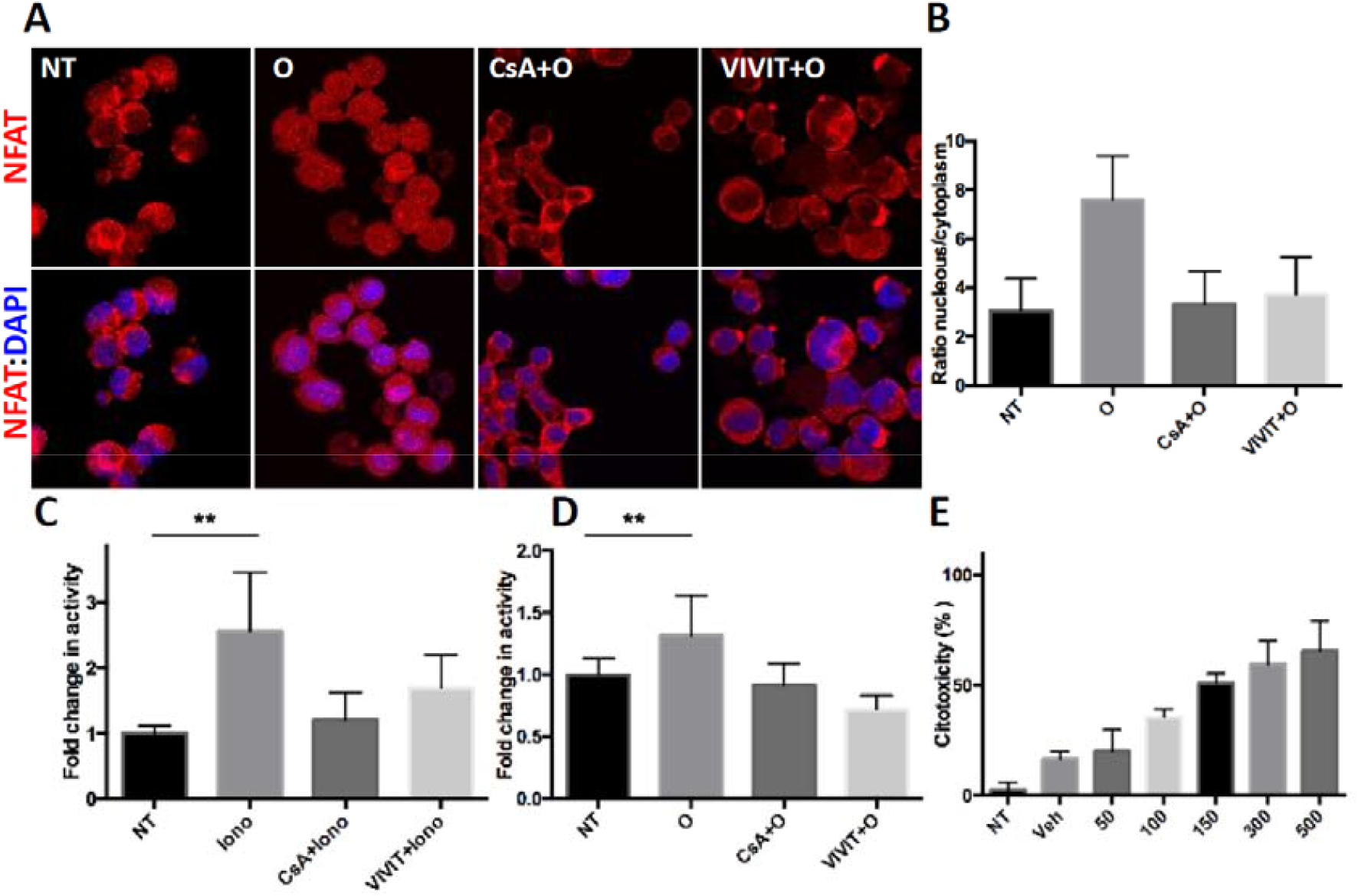
A30P-aSyn-O induce NFAT translocation into the nucleus and cell death in differentiated N2a dopaminergic cells. Differentiated N2a cells were treated with 10 μMA30P-aSyn-O for three hours (**A**). Cells were immunostained with anti-NFATc3 and DAPI and imaged on a confocal microscope. The extent of NFAT translocation into nucleus was calculated by measuring the ratio of the red fluorescence intensity in the nucleus by the red fluorescence intensity in the cytosol (**B**). Differentiated N2a cells were co-transfected with pGL4.30 plasmids containing firefly (reporter gene) and renilla luciferase (control gene). The firefly luciferase gene is under the regulation of 3x NFAT promoter, while renilla luciferase is under the regulation of a constitutive promoter, serving as a normalizing control. N2a cells were treated with ionomycin 3μM(**C**) or A30P-aSyn-O 10 μM (**D**). After 18 h, a dual-luciferase assay was performed to evaluate NFAT activation measured by firefly luciferase luminescence. Cells were treated with 50-500 nM of OHT (Tamoxifen) as indicated in the X axis and after 48 h cytotoxicity was measured by LDH release (**E**). All experiments were performed at least three times and graphs are represented as mean and SD where * p > 0.05, ** p > 0.01 and *** p > 0.001.

To further confirm that A30P-aSyn-O could activate the NFAT pathway, translocating the TF to the nucleus and activating target genes, we used a plasmid encoding for the firefly luciferase reporter under the control of the NFAT promoter (3x NFAT). After transfection with this plasmid, cells were differentiated into DA neurons as described above. As a control, we used that the calcium ionophore ionomycin, a well-known NFAT activator, which caused 2.5-fold increase in the expression of firefly luciferase, confirming that NFAT was activated, migrating to the nucleus, and activating the expression of the reporter gene (**Figure 4C**). As expected, CsA and VIVIT abolished the effect induced by ionomycin, by blocking the NFAT pathway (**Figure 4C**). Importantly, we found that A30P-aSyn-O activated the transcription of the luciferase reporter gene by 1.3-fold, and again, CsA and VIVIT abolished this effect (**Figure 4D**). Taken together, these data show that A30P-aSyn-O induce nuclear translocation and functional activation NFAT, likely mediating cytotoxicity and neuronal death.

To test if NFAT activation was sufficient to induce death of differentiated N2a, cells were transduced with retroviruses encoding CA-NFATc2-ER to stably express a constitutively active NFAT (CA-NFATc2) protein that is activated by tamoxifen (OHT), by fusing the previously characterized CA-NFATc2 to the N-terminally truncated G525R estrogen receptor (ER™), which is only responsive to OHT^48^. The use of a conditioned NFATc2 protein enables temporal control and synchronization of NFAT activation and, therefore, a more detailed analysis of the gene expression profile mediated by this factor and cell death kinetics. Furthermore, this construct bypasses the need for a Ca^2+^ influx to activate NFATc2 protein, reducing the background generated by the induction of other Ca^2+^-responsive transcription factors, such as CREB and MEF2, or of other aSyn-O-induced pathways. As expected, OHT did not induce cell death in differentiated wildtype control N2a cells (data not shown). Interestingly, in differentiated CA-NFATc2-ER N2a expressing cells, OHT induced a marked and concentration dependent cell death (**Figure 4E [OHT]**). Altogether, these data indicate that NFAT transcription factor is activated by A30P-aSyn-O and is sufficient to induce cell death in differentiated N2a neurons, suggesting a main role for this transcription factor in PD pathology.

### Expression of NFAT-regulated genes changes upon treatment with aSyn oligomers

Since NFAT is a transcription factor, and given its involvement in the aSyn-O-induced cytotoxicity and synapse loss in both primary mesencephalic cultures and in differentiated N2a cells, we performed real-time qPCR assays to assess the involvement of the NFAT pathway in regulating gene expression in response to aSyn-O. We assembled a real-time qPCR panel of 91 genes related to various cellular processes involved in PD/synucleinopathies, including inflammation, apoptosis, synaptic modulation, and neural development genes (**Supplementary Table 1**), and assessed gene expression alterations in differentiated N2a treated with A30P-aSyn-O, in the absence or in the presence of VIVIT. Genes with at least 2 folds up- or down-regulation, and that had their expression normalized by VIVIT (**Supplementary Table 1**), were further validated by real-time qPCR in primary neurons and in CA-NFATc2-ER N2a cells (**Figure 5**). Of the 91 genes tested, we identified 5 that were aSyn-O responsive (2-fold up or down-regulated) and modulated by the NFAT pathway, as analyzed by VIVIT inhibition: ***Syn1, Tubb4***, *Kalrn, Bcl-2A1A* and *Bim* (in bold the genes downregulated). We identified 8 genes that were aSyn-O responsive, but not responsive to NFAT inhibition: *Bbc3, Mcl1*, ***Xiap, Pmaip1, Bmp4, Map2K4, Ngf*** and ***Nos2*** (**Supplementary Table 1**). Interestingly the genes *Syn1, Tubb4, Kalrn* are known to be involved in formation and maintenance of synapses, and the other two, *Bcl-2A1A* and *Bim* are known to be involved in apoptosis. Furthermore, we found that the levels of synapsin 1 (Syn1) protein are drastically reduced in primary mesencephalic neurons treated with A30P-aSyn-O, supporting that this gene is regulated by the NFAT transcription factor (**Figure 2**). Furthermore, the genes that were regulated only by oligomers but did not respond to VIVIT NFAT-pathway inhibition, suggest that these genes are likely enrolled in other cellular pathways.

**Figure 5.**
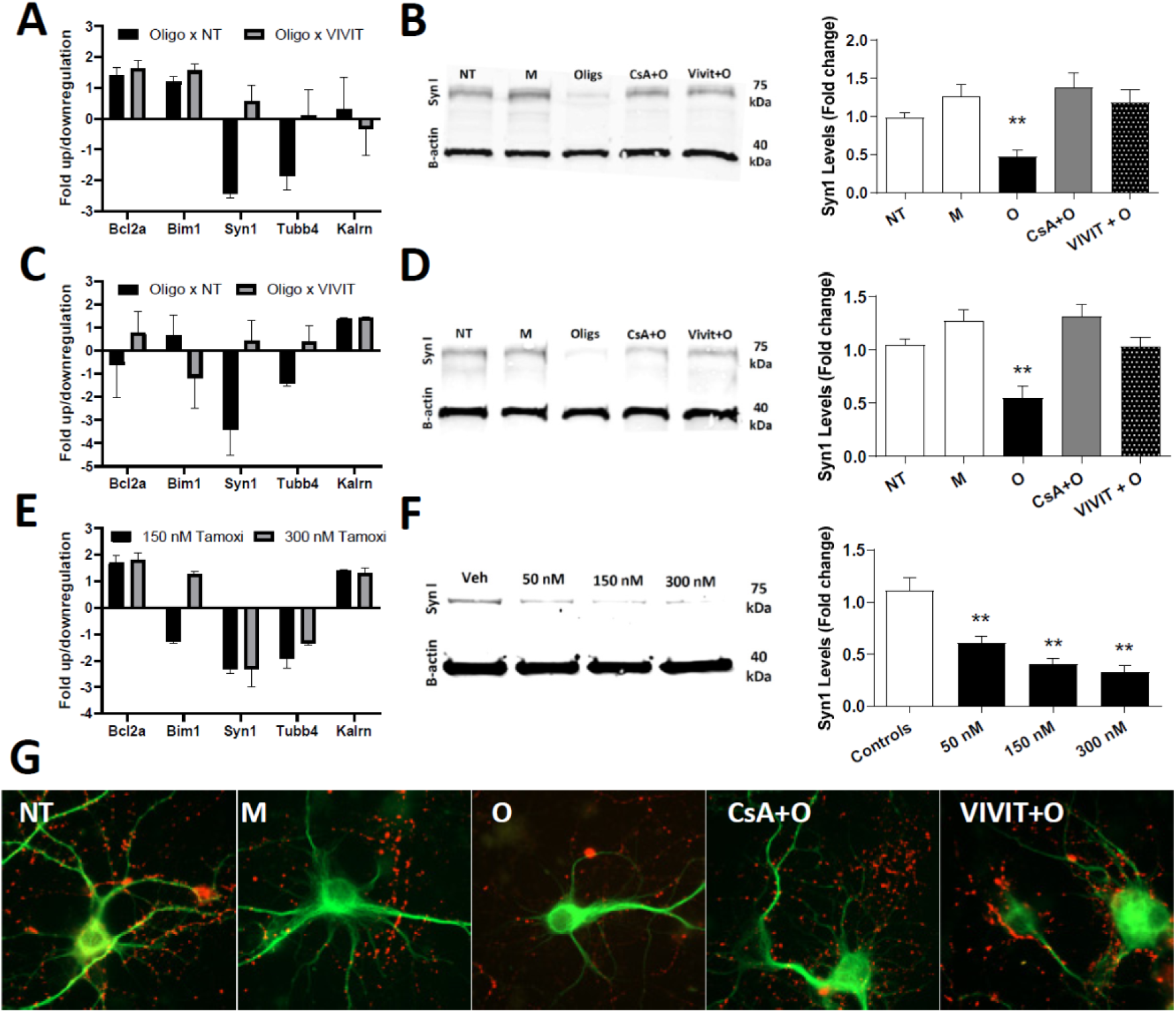
Synapsin 1 levels are decreased in the presence of A30P-aSyn oligomers in an NFAT-dependent pathway. Mouse primary mesencephalic neurons (**A**) or differentiated N2a cells (**C**) were treated with A30P-aSyn-O (10 μM) in the absence or in the presence of VIVIT (10 μM) for 18 h. Genes responsive to NFAT pathway modulation found in the panel (*Syn1, Tubb4, Kalrn, Bcl-2A1A* and *Bim*) were validated by real time PCR. CA-NFAT-ER N2a cells were treated with 150 of OHT (Tamoxifen) and were analyzed after 24 h by real-time-PCR for the same genes (**E**). Western blot analyses were made for the same cells and treatments to assess expression level of Syn1 protein (**B, D and F; left**). β-actin was used as loading control. Graphical representation of band intensities of Syn1 normalized to β-actin (**B, D and F; right**). Mouse primary mesencephalic neurons were treated with 10 μM of A30P-aSyn-O in the absence or presence of 2 μM of CsA or 10 μM of VIVIT for 24 h. Cells were immunostained with anti-MAP2 and anti-synapsin-1 antibodies (**G**). All experiments were performed at least three times and graphs are represented as mean and SD where * p > 0.05, ** p > 0.01 and *** p > 0.001.

To validate the aSyn-O NFAT regulated genes (*Syn1, Tubb4, Kalrn, Bcl-2A1A* and *Bim*), we first repeated the real-time qPCR in primary dopaminergic neurons and in differentiated N2a (**Figure 5A and C)**. Of the five tested only *Syn1* gene showed the same pattern of expression in both models. Surprisingly, NFAT pathway inhibition by VIVIT completely reversed the suppression of this gene induced by aSyn-O.

Furthermore, to explore a possible regulation of these gene exclusively and directly by NFAT activation, CA-NFATc2-ER N2a cells were treated with OHT. In this model, only *Syn1* and *Tubb4* were at least 2-fold down regulated after OHT treatment, indicating that the NFAT transcription factor pathway can directly and sufficiently modulate these genes expression at mRNA level (**Figure 5E**). To our knowledge, this is the first demonstration of a possible direct regulation of the *Syn1* gene by NFAT transcription factor. Consistently, we identified two well conserved putative NFAT-binding sites in the *Syn1* promoter region between five different animal species (**Supplementary Table 2**), whereas no putative NFAT-binding site was found in genes responsive to aSyn-O but irresponsive to VIVIT (genes not regulated by NFAT pathway).

Since the levels of *Syn1* mRNA were the most responsive to A30P-aSyn-O treatment and to NFAT pathway inhibition, we further analyzed Syn1 protein levels in primary neurons and in differentiated N2a cells. Consistent with the reduction in mRNA levels, A30P-aSyn-O significantly reduced Syn1 protein levels (approximately 2-fold) (**Figure 5B and D**). This effect was completely abolished by CsA or VIVIT treatment, strongly suggesting a role of the NFAT pathway in this regulation. Finally, the sole activation of NFAT (in CA-NFATc2-ER N2a by OHT treatment) induced a dose response reduction of Syn1 levels, demonstrating that NFAT can regulate (direct or indirectly) the *Syn1* gene (**Figure 5F**). To further illustrate this regulation, VIVIT treatment also completely reversed the effects of A30P-aSyn-O on presynaptic vesicles, as seen by Syn1 staining in dopaminergic neurons, and restored neuronal morphology to levels comparable to control and CsA treatment (**Figure 5G**). Together, these data demonstrate that A30P-aSyn-O can disrupt neuronal synapses by the activation of NFAT signaling pathway and subsequent downregulation of Syn1 expression suggesting NFAT pathway inhibition might be a target for pharmacological modulation in synucleinopathies.

## Discussion

In this study, we took advantage of the increased oligomerization propensity of a mutant variant of aSyn associated with familial forms of PD (A30P-aSyn) to ascertain the signaling cascades triggered by oligomer-enriched aSyn preparations. We investigated the role of calcineurin-NFAT signaling pathway in the neurotoxicity induced by aSyn oligomeric and fibrillar species. Importantly, we show, for the first time, the direct implication of the NFAT family of transcription factors pathway in the cytotoxicity and synaptic regulation induced by a particular type of extracellular aSyn oligomers. We show that extracellular aSyn oligomers, but not fibrillar species, strongly induce cell death and loss of the pre-synaptic protein synapsin 1, and that these effects are completely reverted by calcineurin (CnA) inhibition with ciclosporin A (CsA) and, surprisingly, also with VIVIT, a specific peptide inhibitor of the CnA-NFAT pathway. This suggests NFAT might be an important transcription factor in PD. Furthermore, NFAT pathway regulation is sufficient, as shown by exclusive NFAT activation (CA-NFATc2-ER N2a), and necessary to induce PD-related cellular phenotypes, including neural cell death, possibly by regulating important genes involved in apoptosis and synapsin 1 expression.

aSyn overexpression, extracellular accumulation, or administration, have been used to generate numerous cell and animal models of PD, as this protein is considered a major culprit in PD ^57^. As previously shown, oligomeric WT or mutant aSyn induce cell death in dopaminergic cell lines and in primary neuronal cultures, alter functional neuronal markers, decrease synaptic plasticity, and induce memory deficits^37,57,58^. Cells must tightly regulate Ca^2+^ homeostasis to prevent pathological disorders and cell death due to excitotoxicity^59^. Extracellular oligomeric forms of aSyn induce Ca^2+^ overload into neurons, and the consequent activation of downstream pathways leads to pathological alterations and, ultimately, to cell death and several other PD-related alterations^37,39,60–63^. One hypothesis is that aSyn certain oligomers may induce the formation of Ca^2+^-permeable pores at the plasma membrane, in accord to what has been postulated for other aggregating proteins, such as amyloid-β peptides and prion proteins^57,64^.

One of the best-characterized pathways activated by Ca^+2^ influx is the Ca^2+-^ CaM-dependent serine/threonine phosphatase calcineurin, an essential enzyme that plays a key role in neurite extension, synaptic plasticity, memory, and learning^65^, and is implicated as a key mediator of aSyn toxicity. Pharmacological inhibition of the catalytic subunit of calcineurin, calcineurin A (CnA), with clinically used inhibitors, such as FK506 and CsA, can reduce aSyn toxicity in different models^37,38,66–68^. Although the role of Ca^2+^-CaM-CnA pathway in aSyn induced phenotypes has been documented, the downstream signaling pathways are still obscure. One of the main transcription factors targeted by CnA dephosphorylation is the NFAT-family^44^. Here, we showed that A30P-aSyn-O can activate the NFAT transcription factor leading to its translocation into the nucleus and activation of reporter a gene (**Figure 4**). Moreover, we showed that inhibition of CnA with CsA and with VIVIT, a specific inhibitor of the CnA-NFAT pathway, can revert this transcription factor activation and block aSyn-O neurotoxicity, as characterized by the almost complete inhibition of cell death, reversal of neural ramification deterioration, and loss of pre-synaptic synapsin 1 positive puncta formation (**Figures 1, 2 and 3**). Consistently, the NFAT transcription factor has already been demonstrated to be important in neural and axonal development^47^, survival^69^, and death^70–72^, being localized in the nucleus of neurons in aSyn transgenic mice in postmortem brain tissue from PD patients^38^.

However, the direct role of this transcription factor in neuronal degeneration and the regulation of the expression of relevant PD-related genes has not been determined. After the analysis of a panel of 91 genes related to various cellular processes involved in PD/synucleinopathies (including inflammation, apoptosis, synaptic modulation, and neural development) by real-time qPCR, we found several putative genes regulated by NFAT induced by A30P-aSyn-O. In differentiated N2a cells, *Bcl2a* and *Bim*, two apoptotic related proteins, seemed to be regulated by NFAT. However, after validation in primary neurons and in N2a cells with only the NFAT pathway specifically activated (NFAT-ER), we found their expression was close to the detection limit of the technique, and that their pattern varied between different cell types. NFAT directly regulates the expression of several apoptosis-related genes such as *Fas-L, Nur77, c-Flip* and *TNF-*α, *A1* and *Bcl2* in several other cell types^42,49,73^. In the future, further analysis will be needed to unveil how NFAT pathway regulates neuron cell death induced by aSyn-O, since both its inhibition and exclusive activation is sufficient to modulate this process.

Interestingly, one gene that might be directly regulated by NFAT at the transcriptional and translational level is *Syn1*, that encodes synapsin 1, as this was validated in our three different cells models (**Figure 5**). Several recent studies have suggested that intra- or extracellular aSyn overexpression or accumulation can induce presynaptic dysfunction by disturbing vesicular trafficking, leading to defects in neurotransmitter release ^74–78^ and to the loss of several presynaptic proteins important for neurotransmitter release and synapse formation^54,74,79^. Synapsin 1 is of major importance because of its role in vesicle trafficking. In resting conditions, synapsin 1 attaches the vesicle to the actin-based cytoskeleton. During excitation, and upon phosphorylated by different kinases, it detaches from the vesicles facilitating neurotransmitter release and mediates the binding of recycling vesicles to the actin cytoskeleton^80–82^. Interestingly, aSyn aggregation has been suggested to down-regulate the expression of synapsin 1, correlating with PD phenotypes such as locomotion and memory impairment^54,74,79,83^. A recent study also suggested that synapsins are implicated in the inhibition of synaptic vesicle exocytosis by aSyn^84^. Finally, at a molecular level, the transcriptional expression of synapsin genes (*Syn1*) is down-regulated *in vivo* and *in vitro* by aSyn oligomers^54^. However, although synapsin 1 was already described as an important target for oligomeric aSyn, the molecular mechanisms involved in the transcriptional regulation of synapsin 1 by aSyn were unknown. CnA inhibition has been shown to reverse neuritic beading and rescue structural disruption of neuronal network in the presence of Aβ^85^, and to regulate different proteins involved in neurotransmitter release, including synapsins, synaptotagmin, rabphilin2A, synaptobrevin, and dephosphins^86^. However, we show here for the first time, that NFAT transcription factor has two possible highly conserved putative binding sites in the *Syn1* promoter, and is necessary and sufficient for regulating *Syn1*, leading to its downregulation at the transcriptional (mRNA) and translational level (protein). NFAT inhibition leads to synapsin 1 expression and accumulation at presynaptic puncta in neurons, where it performs its normal function. NFAT can bind to specific DNA-responsive elements defined as (A/T)GGAAA(A/N) (A/T/C) to activate or deactivate gene transcription, either alone or in synergy with several other transcriptional regulators^42,87^. Interestingly, we identified two well-conserved (at least in five different animal species) putative NFAT-responsive elements in the *Syn1* promoter region, adding more evidence supporting NFTA regulation of synapsin 1 gene translation.

Therefore, our current results reveal a previously undocumented mechanism whereby small oligomeric aSyn species may disturb synapses in PD, possibly affecting memory function and cognitive ability (**Figure 6**). Our results further suggest that such pathological mechanisms may be effectively targeted by pharmacological inhibition of NFAT by selective small-molecule inhibitors of calcineurin-NFAT signaling that mimic VIVIT, or by already-approved drugs as CsA and FK506.

**Figure 6.**
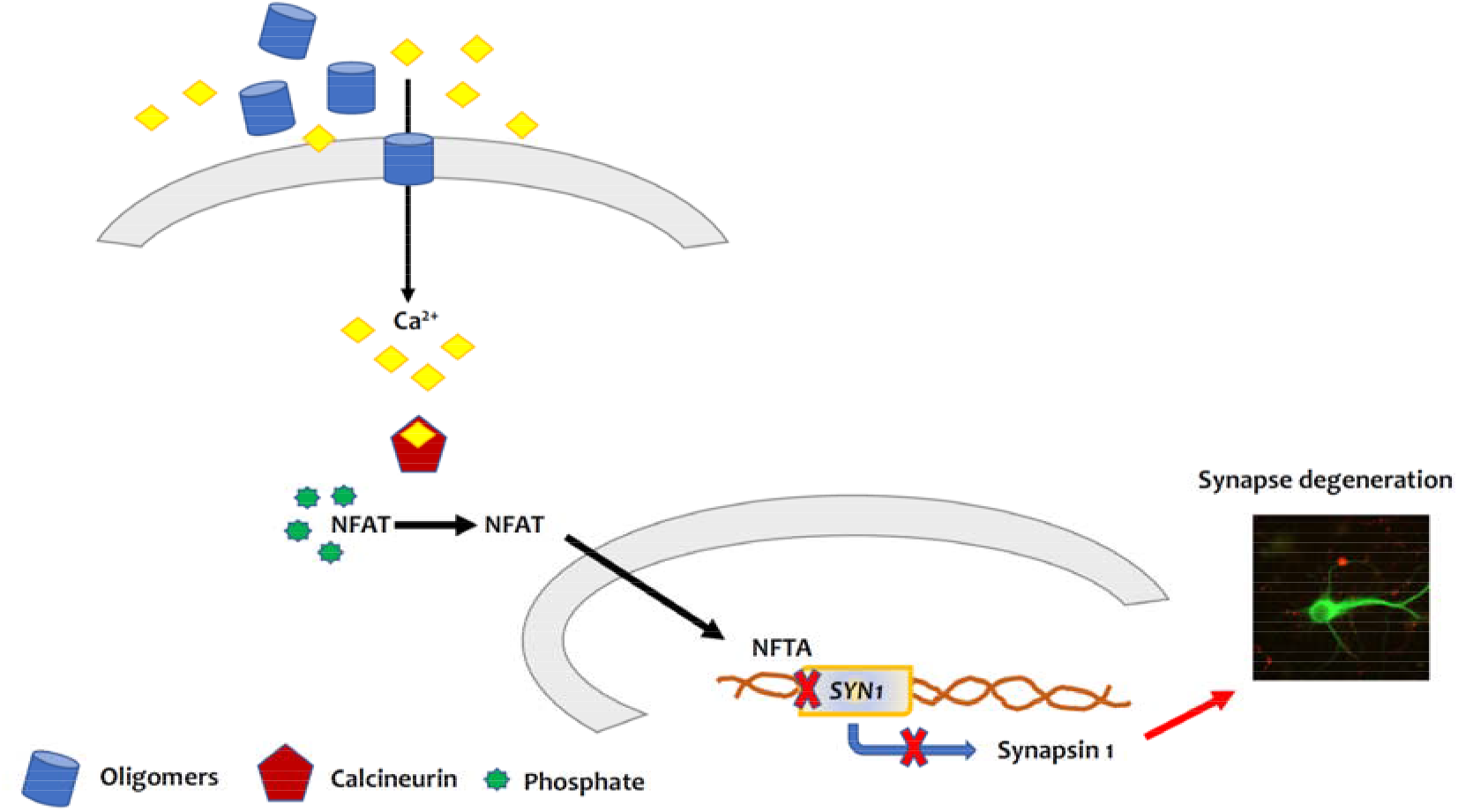
Schematic representation of the participation of NFAT on synapse degeneration in synucleinopathies. A specific population of aSyn-O present in the extracellular milieu could be inserted in the plasma membrane leading to leakage of calcium to the interior of the neurons. Upon calcium binding, activation of calcineurin takes place. This phosphatase dephosphorylates the hyperphosphorylated, inactive NFAT leading to its activation and translocation to the nucleus. There, NFAT binds to its cognate promoter sequences including the one that regulates the expression of the gene of synapsin 1 (*Syn1*). NFAT downregulates the expression of this gene decreasing the amount synapsin 1, a protein that is involved in vesicle trafficking facilitating neurotransmitter release. This cascade of events leads to synapse degeneration one of the events observed in PD and other synucleionopathies.

## Acknowledgements

TFO is supported by the Deutsche Forschungsgemeinschaft (DFG, German Research Foundation) under Germany’s Excellence Strategy - EXC 2067/1-390729940, and by SFB1286 (B8). DF is supported by the Federal Brazilian Funding Agencies, CNPq and Capes, and by the State of Rio de Janeiro Funding Agency, Faperj. RS is supported by the Federal Brazilian Funding Agencies, CNPq and Capes, and by the State of Rio de Janeiro Funding Agency, Faperj.

## Figure legends

**Supplementary figure 1.**
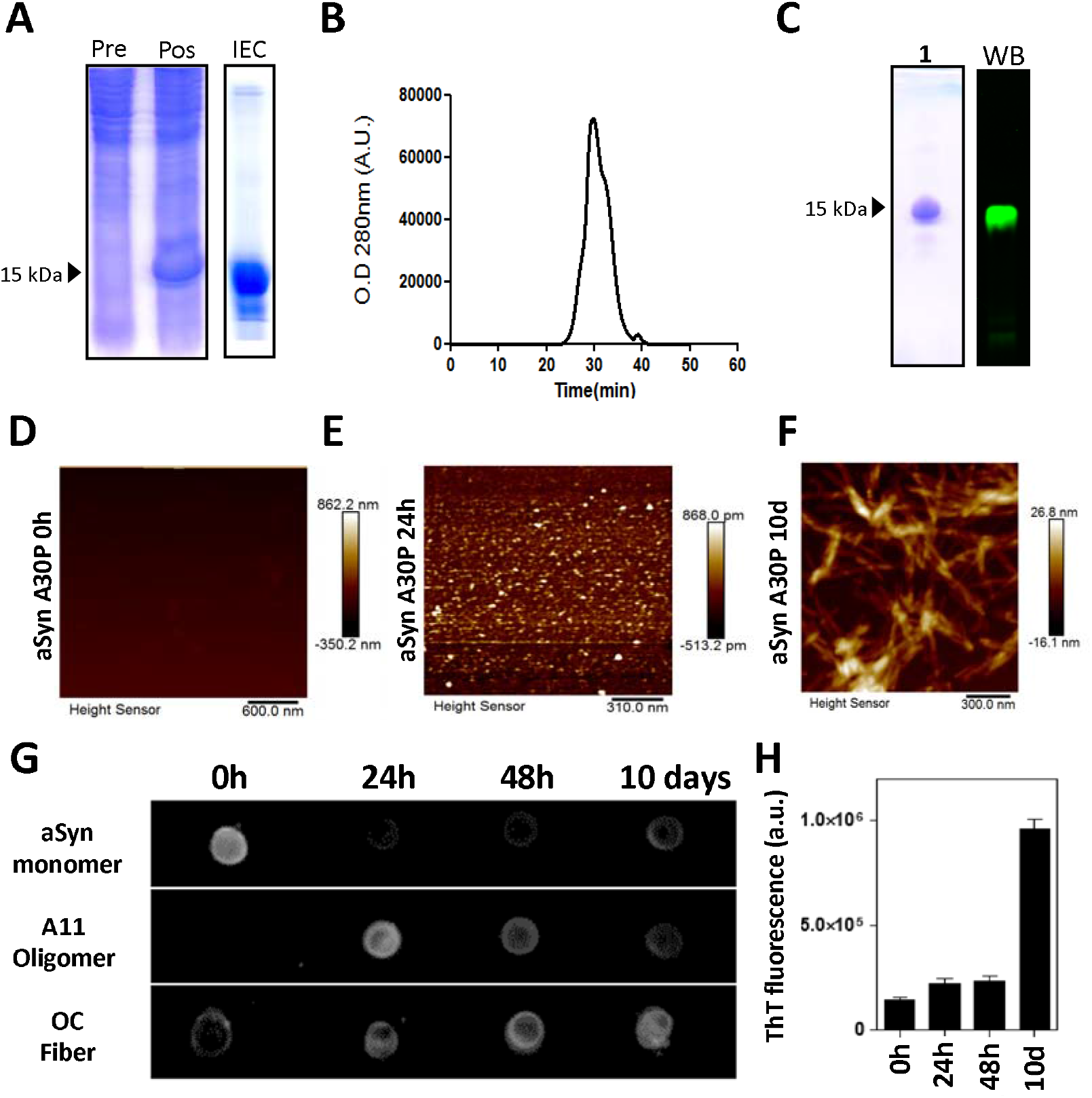

**Supplementary Table 1.**
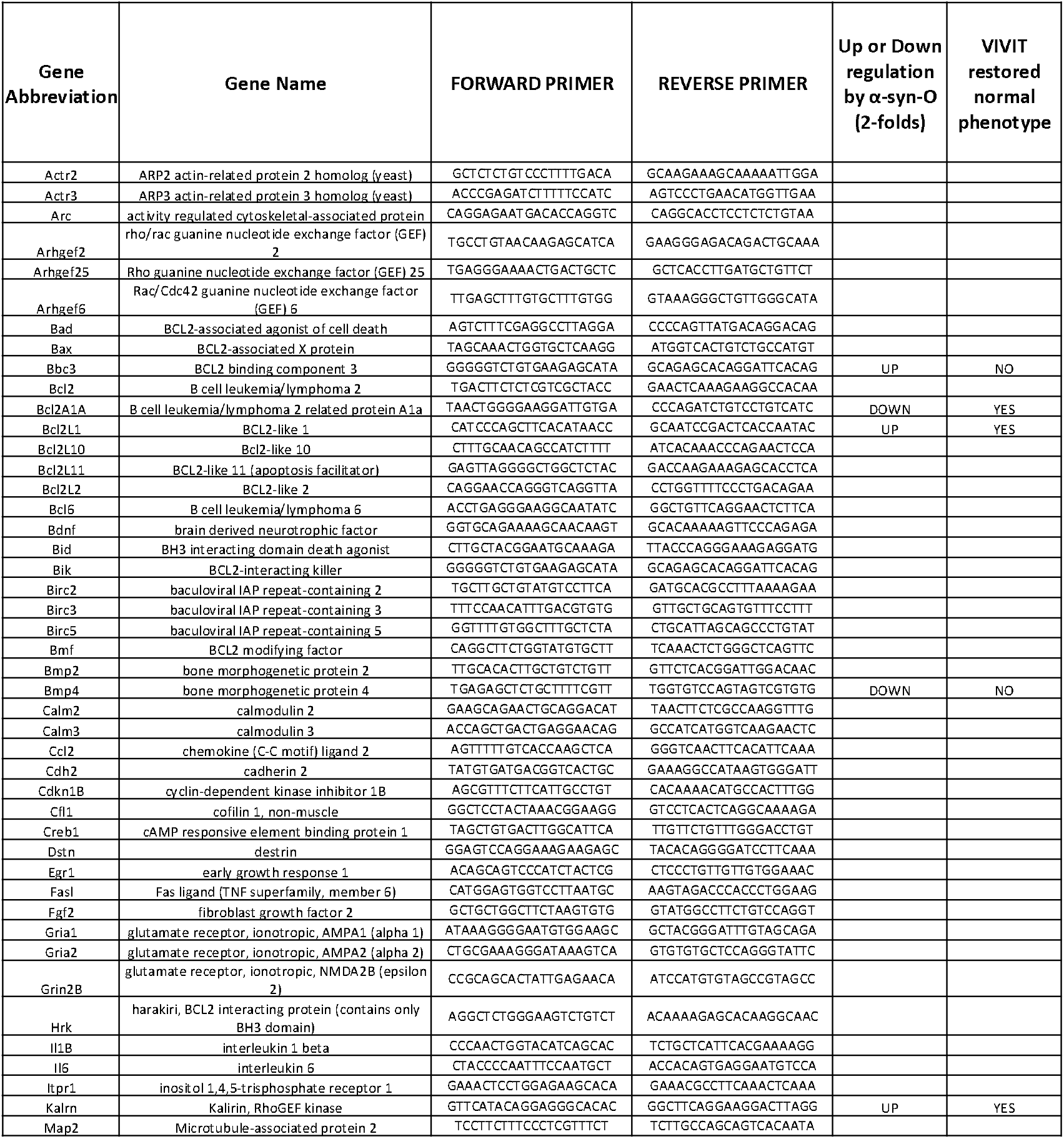
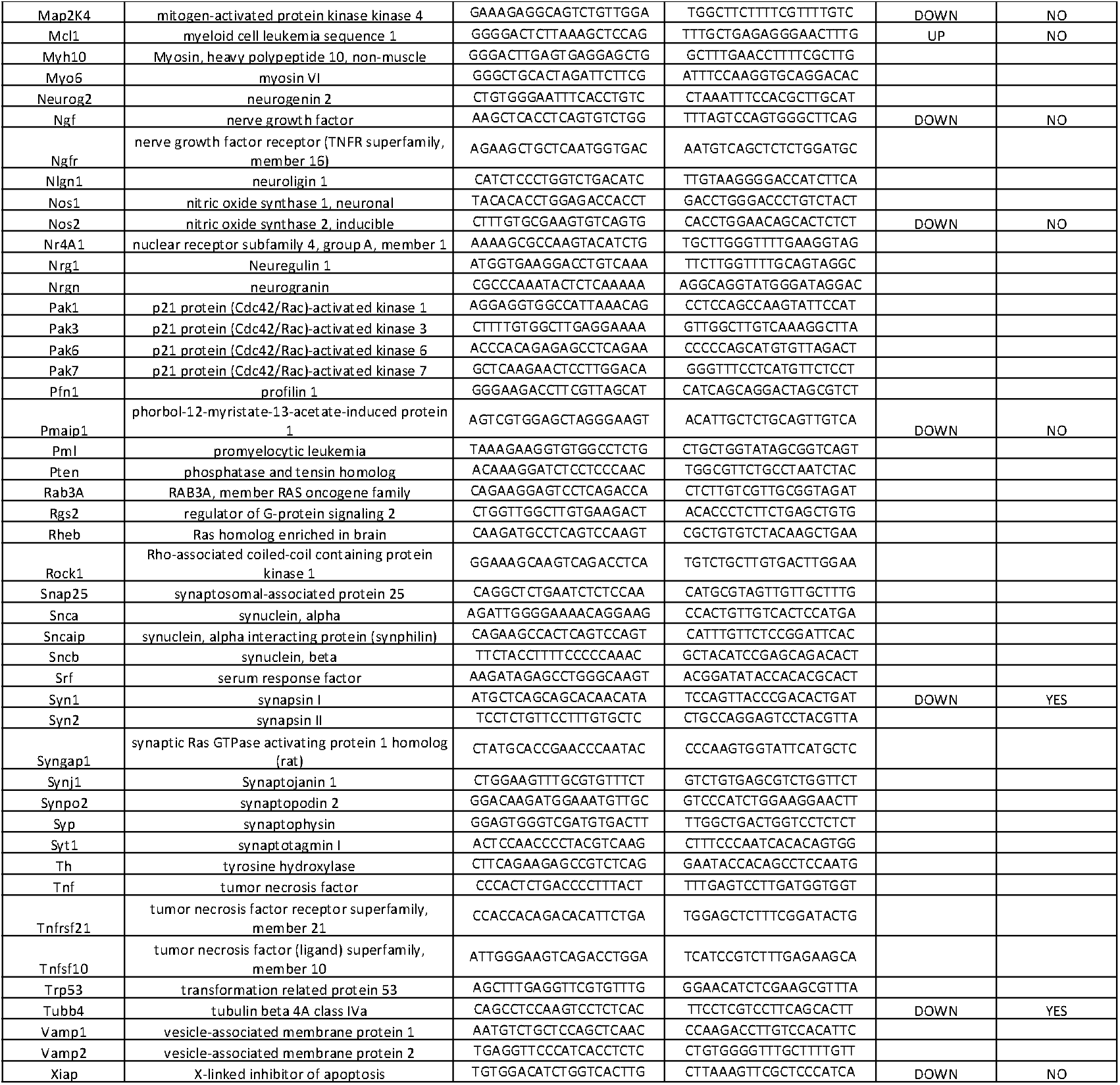
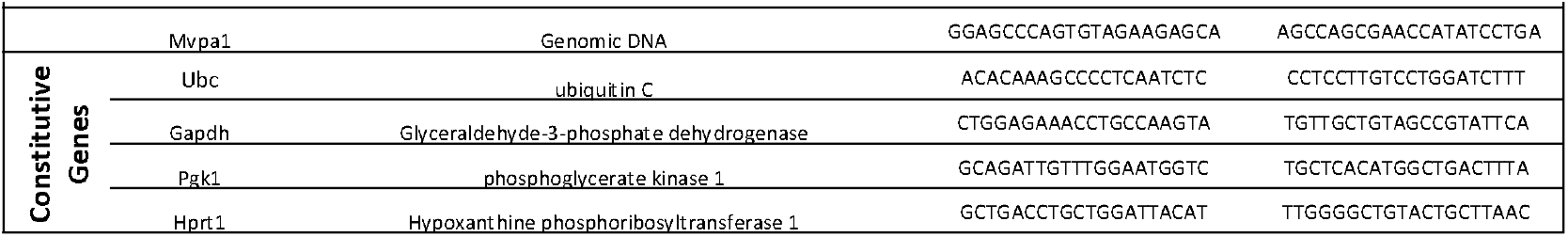

**Supplementary Table 2.**
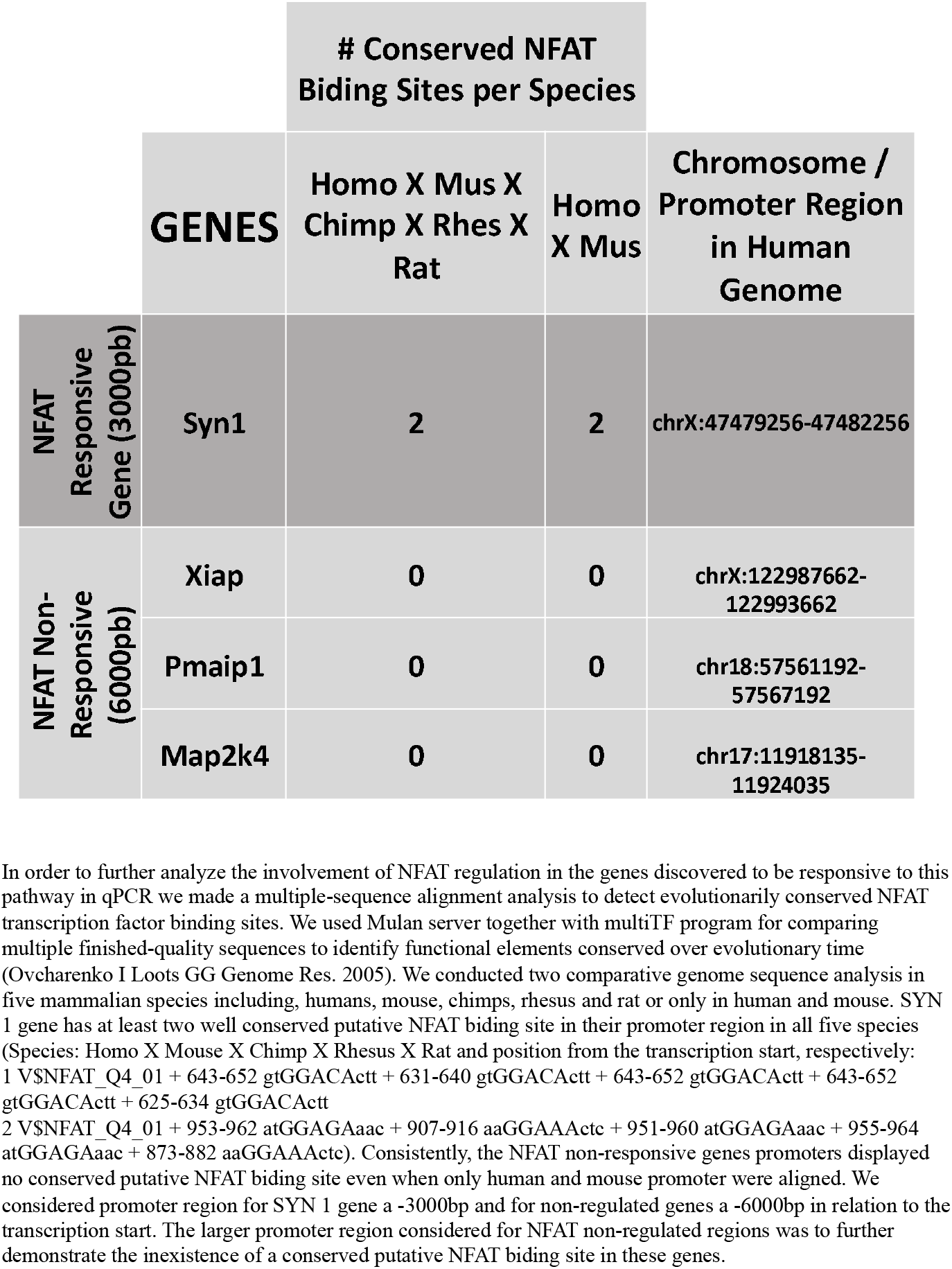

## Notes

### Competing Interest Statement

The authors have declared no competing interest.

### Summary of Updates

Important reference needed to be added

