## Supplementary Figure 1 for "Alpha-synuclein oligomers activate NFAT proteins modulating synaptic homeostasis and apoptosis"

### Slide 1
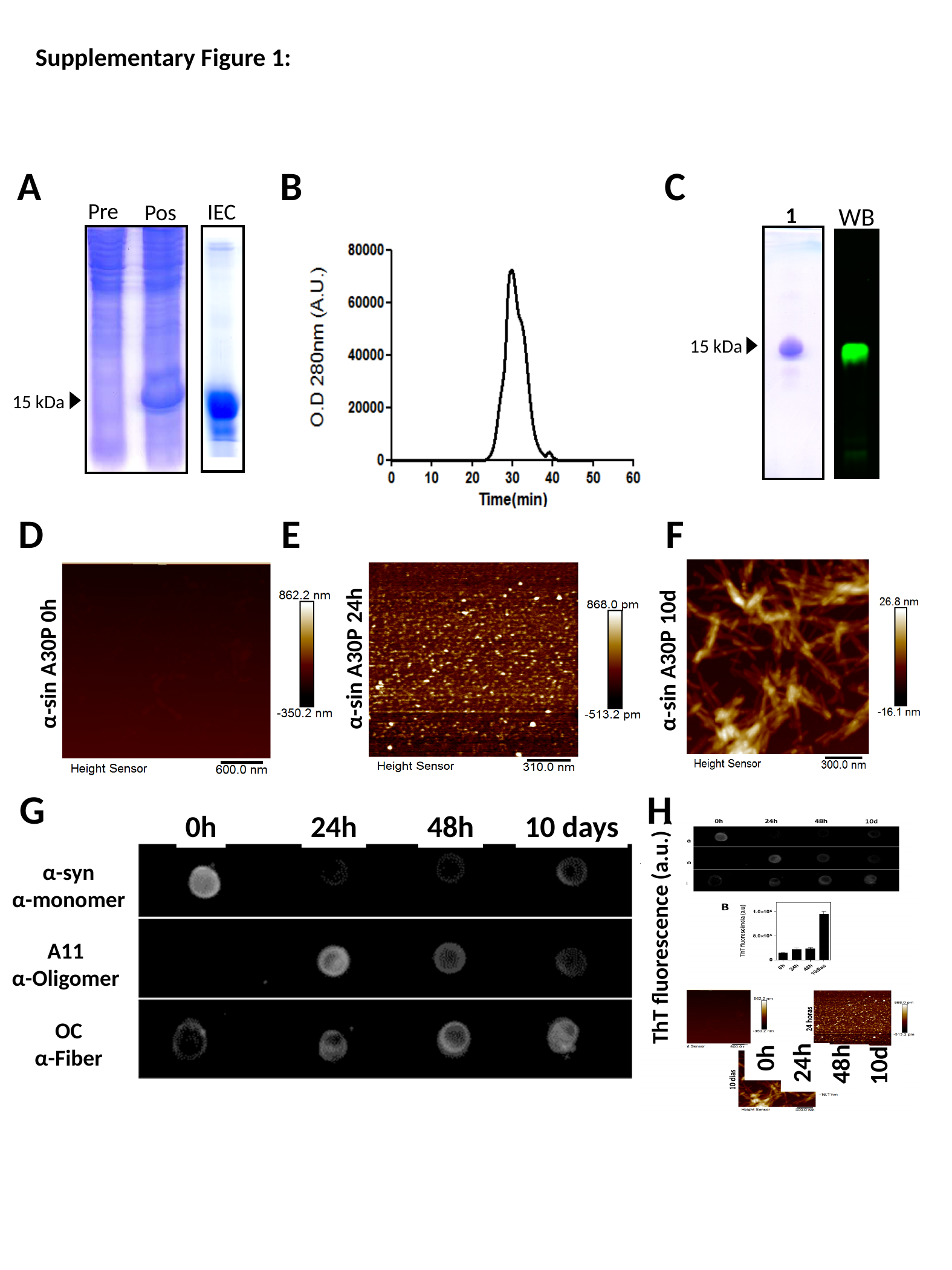

Supplementary Figure 1:
A
B
C
Pre
IEC
Pos
WB
1
15 kDa
15 kDa
D
E
F
α-sin A30P 24h
α-sin A30P 0h
α-sin A30P 10d
G
H
0h
24h
48h
10 days
ThT fluorescence (a.u.)
24h
48h
0h
10d
α-syn
α-monomer
A11
α-Oligomer
OC
α-Fiber

### Slide 2
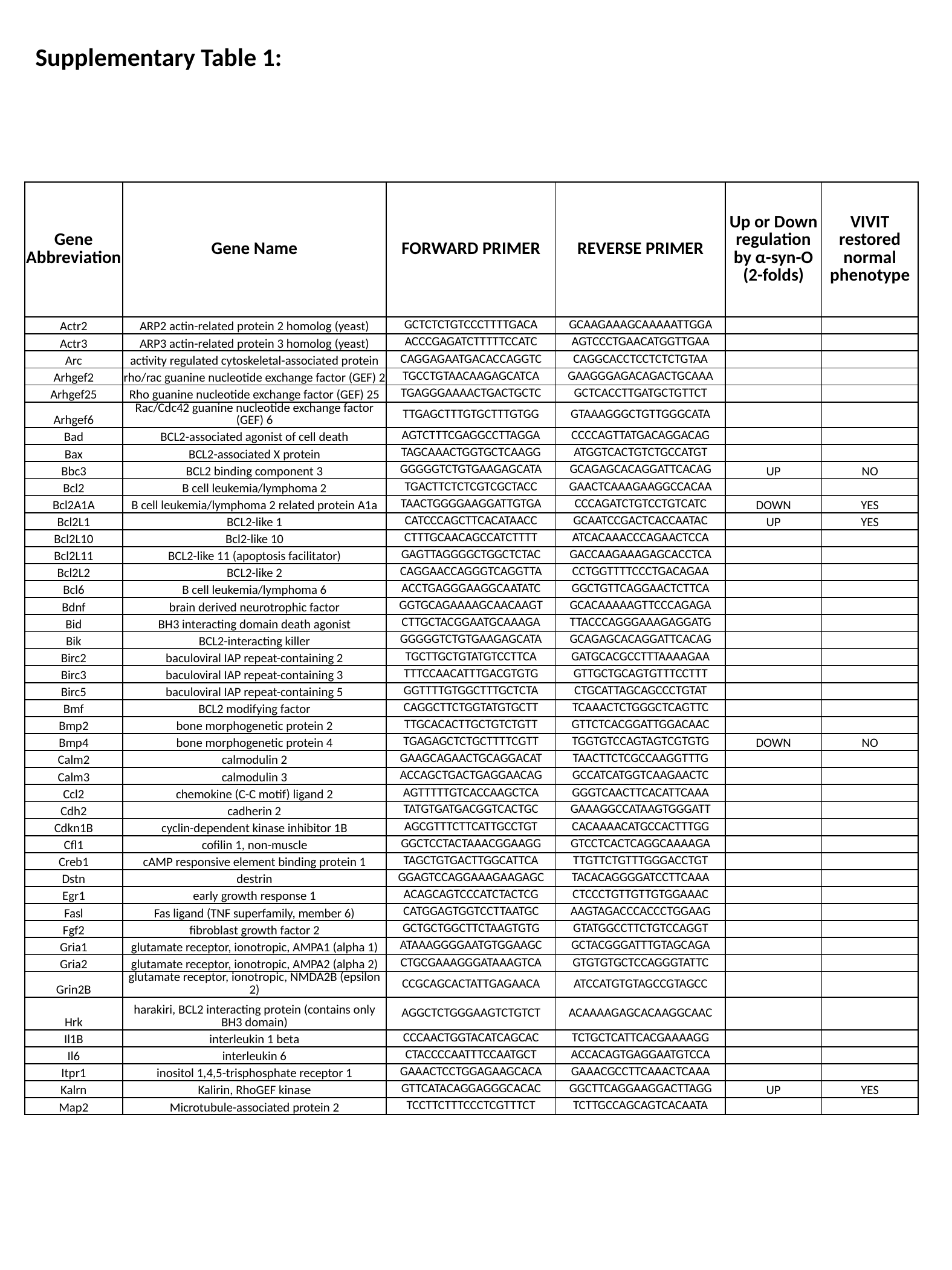

Supplementary Table 1:
| Gene Abbreviation | Gene Name | FORWARD PRIMER | REVERSE PRIMER | Up or Down regulation by α-syn-O (2-folds) | VIVIT restored normal phenotype |
| --- | --- | --- | --- | --- | --- |
| Actr2 | ARP2 actin-related protein 2 homolog (yeast) | GCTCTCTGTCCCTTTTGACA | GCAAGAAAGCAAAAATTGGA | | |
| Actr3 | ARP3 actin-related protein 3 homolog (yeast) | ACCCGAGATCTTTTTCCATC | AGTCCCTGAACATGGTTGAA | | |
| Arc | activity regulated cytoskeletal-associated protein | CAGGAGAATGACACCAGGTC | CAGGCACCTCCTCTCTGTAA | | |
| Arhgef2 | rho/rac guanine nucleotide exchange factor (GEF) 2 | TGCCTGTAACAAGAGCATCA | GAAGGGAGACAGACTGCAAA | | |
| Arhgef25 | Rho guanine nucleotide exchange factor (GEF) 25 | TGAGGGAAAACTGACTGCTC | GCTCACCTTGATGCTGTTCT | | |
| Arhgef6 | Rac/Cdc42 guanine nucleotide exchange factor (GEF) 6 | TTGAGCTTTGTGCTTTGTGG | GTAAAGGGCTGTTGGGCATA | | |
| Bad | BCL2-associated agonist of cell death | AGTCTTTCGAGGCCTTAGGA | CCCCAGTTATGACAGGACAG | | |
| Bax | BCL2-associated X protein | TAGCAAACTGGTGCTCAAGG | ATGGTCACTGTCTGCCATGT | | |
| Bbc3 | BCL2 binding component 3 | GGGGGTCTGTGAAGAGCATA | GCAGAGCACAGGATTCACAG | UP | NO |
| Bcl2 | B cell leukemia/lymphoma 2 | TGACTTCTCTCGTCGCTACC | GAACTCAAAGAAGGCCACAA | | |
| Bcl2A1A | B cell leukemia/lymphoma 2 related protein A1a | TAACTGGGGAAGGATTGTGA | CCCAGATCTGTCCTGTCATC | DOWN | YES |
| Bcl2L1 | BCL2-like 1 | CATCCCAGCTTCACATAACC | GCAATCCGACTCACCAATAC | UP | YES |
| Bcl2L10 | Bcl2-like 10 | CTTTGCAACAGCCATCTTTT | ATCACAAACCCAGAACTCCA | | |
| Bcl2L11 | BCL2-like 11 (apoptosis facilitator) | GAGTTAGGGGCTGGCTCTAC | GACCAAGAAAGAGCACCTCA | | |
| Bcl2L2 | BCL2-like 2 | CAGGAACCAGGGTCAGGTTA | CCTGGTTTTCCCTGACAGAA | | |
| Bcl6 | B cell leukemia/lymphoma 6 | ACCTGAGGGAAGGCAATATC | GGCTGTTCAGGAACTCTTCA | | |
| Bdnf | brain derived neurotrophic factor | GGTGCAGAAAAGCAACAAGT | GCACAAAAAGTTCCCAGAGA | | |
| Bid | BH3 interacting domain death agonist | CTTGCTACGGAATGCAAAGA | TTACCCAGGGAAAGAGGATG | | |
| Bik | BCL2-interacting killer | GGGGGTCTGTGAAGAGCATA | GCAGAGCACAGGATTCACAG | | |
| Birc2 | baculoviral IAP repeat-containing 2 | TGCTTGCTGTATGTCCTTCA | GATGCACGCCTTTAAAAGAA | | |
| Birc3 | baculoviral IAP repeat-containing 3 | TTTCCAACATTTGACGTGTG | GTTGCTGCAGTGTTTCCTTT | | |
| Birc5 | baculoviral IAP repeat-containing 5 | GGTTTTGTGGCTTTGCTCTA | CTGCATTAGCAGCCCTGTAT | | |
| Bmf | BCL2 modifying factor | CAGGCTTCTGGTATGTGCTT | TCAAACTCTGGGCTCAGTTC | | |
| Bmp2 | bone morphogenetic protein 2 | TTGCACACTTGCTGTCTGTT | GTTCTCACGGATTGGACAAC | | |
| Bmp4 | bone morphogenetic protein 4 | TGAGAGCTCTGCTTTTCGTT | TGGTGTCCAGTAGTCGTGTG | DOWN | NO |
| Calm2 | calmodulin 2 | GAAGCAGAACTGCAGGACAT | TAACTTCTCGCCAAGGTTTG | | |
| Calm3 | calmodulin 3 | ACCAGCTGACTGAGGAACAG | GCCATCATGGTCAAGAACTC | | |
| Ccl2 | chemokine (C-C motif) ligand 2 | AGTTTTTGTCACCAAGCTCA | GGGTCAACTTCACATTCAAA | | |
| Cdh2 | cadherin 2 | TATGTGATGACGGTCACTGC | GAAAGGCCATAAGTGGGATT | | |
| Cdkn1B | cyclin-dependent kinase inhibitor 1B | AGCGTTTCTTCATTGCCTGT | CACAAAACATGCCACTTTGG | | |
| Cfl1 | cofilin 1, non-muscle | GGCTCCTACTAAACGGAAGG | GTCCTCACTCAGGCAAAAGA | | |
| Creb1 | cAMP responsive element binding protein 1 | TAGCTGTGACTTGGCATTCA | TTGTTCTGTTTGGGACCTGT | | |
| Dstn | destrin | GGAGTCCAGGAAAGAAGAGC | TACACAGGGGATCCTTCAAA | | |
| Egr1 | early growth response 1 | ACAGCAGTCCCATCTACTCG | CTCCCTGTTGTTGTGGAAAC | | |
| Fasl | Fas ligand (TNF superfamily, member 6) | CATGGAGTGGTCCTTAATGC | AAGTAGACCCACCCTGGAAG | | |
| Fgf2 | fibroblast growth factor 2 | GCTGCTGGCTTCTAAGTGTG | GTATGGCCTTCTGTCCAGGT | | |
| Gria1 | glutamate receptor, ionotropic, AMPA1 (alpha 1) | ATAAAGGGGAATGTGGAAGC | GCTACGGGATTTGTAGCAGA | | |
| Gria2 | glutamate receptor, ionotropic, AMPA2 (alpha 2) | CTGCGAAAGGGATAAAGTCA | GTGTGTGCTCCAGGGTATTC | | |
| Grin2B | glutamate receptor, ionotropic, NMDA2B (epsilon 2) | CCGCAGCACTATTGAGAACA | ATCCATGTGTAGCCGTAGCC | | |
| Hrk | harakiri, BCL2 interacting protein (contains only BH3 domain) | AGGCTCTGGGAAGTCTGTCT | ACAAAAGAGCACAAGGCAAC | | |
| Il1B | interleukin 1 beta | CCCAACTGGTACATCAGCAC | TCTGCTCATTCACGAAAAGG | | |
| Il6 | interleukin 6 | CTACCCCAATTTCCAATGCT | ACCACAGTGAGGAATGTCCA | | |
| Itpr1 | inositol 1,4,5-trisphosphate receptor 1 | GAAACTCCTGGAGAAGCACA | GAAACGCCTTCAAACTCAAA | | |
| Kalrn | Kalirin, RhoGEF kinase | GTTCATACAGGAGGGCACAC | GGCTTCAGGAAGGACTTAGG | UP | YES |
| Map2 | Microtubule-associated protein 2 | TCCTTCTTTCCCTCGTTTCT | TCTTGCCAGCAGTCACAATA | | |

### Slide 3
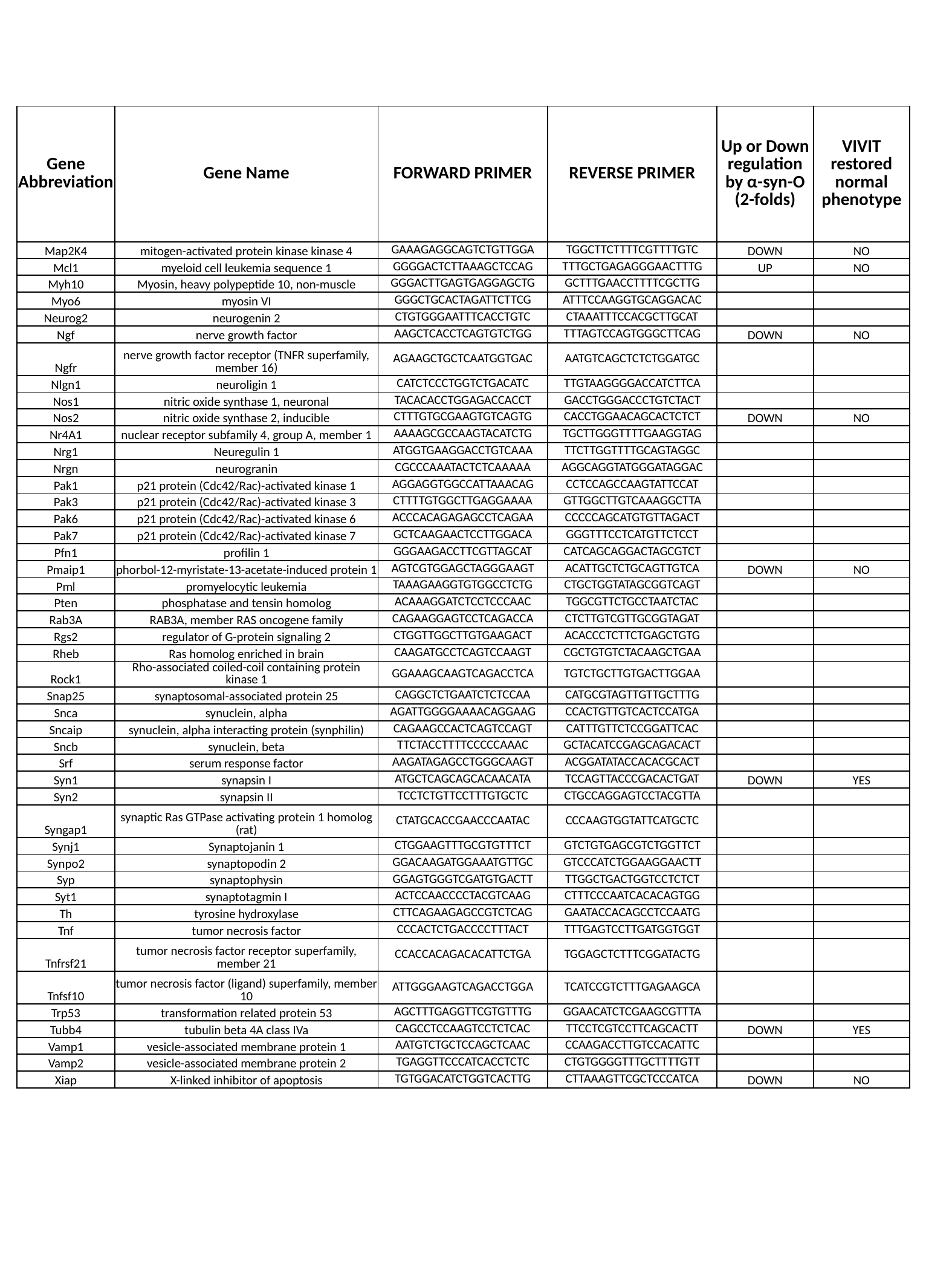

| Gene Abbreviation | Gene Name | FORWARD PRIMER | REVERSE PRIMER | Up or Down regulation by α-syn-O (2-folds) | VIVIT restored normal phenotype |
| --- | --- | --- | --- | --- | --- |
| Map2K4 | mitogen-activated protein kinase kinase 4 | GAAAGAGGCAGTCTGTTGGA | TGGCTTCTTTTCGTTTTGTC | DOWN | NO |
| Mcl1 | myeloid cell leukemia sequence 1 | GGGGACTCTTAAAGCTCCAG | TTTGCTGAGAGGGAACTTTG | UP | NO |
| Myh10 | Myosin, heavy polypeptide 10, non-muscle | GGGACTTGAGTGAGGAGCTG | GCTTTGAACCTTTTCGCTTG | | |
| Myo6 | myosin VI | GGGCTGCACTAGATTCTTCG | ATTTCCAAGGTGCAGGACAC | | |
| Neurog2 | neurogenin 2 | CTGTGGGAATTTCACCTGTC | CTAAATTTCCACGCTTGCAT | | |
| Ngf | nerve growth factor | AAGCTCACCTCAGTGTCTGG | TTTAGTCCAGTGGGCTTCAG | DOWN | NO |
| Ngfr | nerve growth factor receptor (TNFR superfamily, member 16) | AGAAGCTGCTCAATGGTGAC | AATGTCAGCTCTCTGGATGC | | |
| Nlgn1 | neuroligin 1 | CATCTCCCTGGTCTGACATC | TTGTAAGGGGACCATCTTCA | | |
| Nos1 | nitric oxide synthase 1, neuronal | TACACACCTGGAGACCACCT | GACCTGGGACCCTGTCTACT | | |
| Nos2 | nitric oxide synthase 2, inducible | CTTTGTGCGAAGTGTCAGTG | CACCTGGAACAGCACTCTCT | DOWN | NO |
| Nr4A1 | nuclear receptor subfamily 4, group A, member 1 | AAAAGCGCCAAGTACATCTG | TGCTTGGGTTTTGAAGGTAG | | |
| Nrg1 | Neuregulin 1 | ATGGTGAAGGACCTGTCAAA | TTCTTGGTTTTGCAGTAGGC | | |
| Nrgn | neurogranin | CGCCCAAATACTCTCAAAAA | AGGCAGGTATGGGATAGGAC | | |
| Pak1 | p21 protein (Cdc42/Rac)-activated kinase 1 | AGGAGGTGGCCATTAAACAG | CCTCCAGCCAAGTATTCCAT | | |
| Pak3 | p21 protein (Cdc42/Rac)-activated kinase 3 | CTTTTGTGGCTTGAGGAAAA | GTTGGCTTGTCAAAGGCTTA | | |
| Pak6 | p21 protein (Cdc42/Rac)-activated kinase 6 | ACCCACAGAGAGCCTCAGAA | CCCCCAGCATGTGTTAGACT | | |
| Pak7 | p21 protein (Cdc42/Rac)-activated kinase 7 | GCTCAAGAACTCCTTGGACA | GGGTTTCCTCATGTTCTCCT | | |
| Pfn1 | profilin 1 | GGGAAGACCTTCGTTAGCAT | CATCAGCAGGACTAGCGTCT | | |
| Pmaip1 | phorbol-12-myristate-13-acetate-induced protein 1 | AGTCGTGGAGCTAGGGAAGT | ACATTGCTCTGCAGTTGTCA | DOWN | NO |
| Pml | promyelocytic leukemia | TAAAGAAGGTGTGGCCTCTG | CTGCTGGTATAGCGGTCAGT | | |
| Pten | phosphatase and tensin homolog | ACAAAGGATCTCCTCCCAAC | TGGCGTTCTGCCTAATCTAC | | |
| Rab3A | RAB3A, member RAS oncogene family | CAGAAGGAGTCCTCAGACCA | CTCTTGTCGTTGCGGTAGAT | | |
| Rgs2 | regulator of G-protein signaling 2 | CTGGTTGGCTTGTGAAGACT | ACACCCTCTTCTGAGCTGTG | | |
| Rheb | Ras homolog enriched in brain | CAAGATGCCTCAGTCCAAGT | CGCTGTGTCTACAAGCTGAA | | |
| Rock1 | Rho-associated coiled-coil containing protein kinase 1 | GGAAAGCAAGTCAGACCTCA | TGTCTGCTTGTGACTTGGAA | | |
| Snap25 | synaptosomal-associated protein 25 | CAGGCTCTGAATCTCTCCAA | CATGCGTAGTTGTTGCTTTG | | |
| Snca | synuclein, alpha | AGATTGGGGAAAACAGGAAG | CCACTGTTGTCACTCCATGA | | |
| Sncaip | synuclein, alpha interacting protein (synphilin) | CAGAAGCCACTCAGTCCAGT | CATTTGTTCTCCGGATTCAC | | |
| Sncb | synuclein, beta | TTCTACCTTTTCCCCCAAAC | GCTACATCCGAGCAGACACT | | |
| Srf | serum response factor | AAGATAGAGCCTGGGCAAGT | ACGGATATACCACACGCACT | | |
| Syn1 | synapsin I | ATGCTCAGCAGCACAACATA | TCCAGTTACCCGACACTGAT | DOWN | YES |
| Syn2 | synapsin II | TCCTCTGTTCCTTTGTGCTC | CTGCCAGGAGTCCTACGTTA | | |
| Syngap1 | synaptic Ras GTPase activating protein 1 homolog (rat) | CTATGCACCGAACCCAATAC | CCCAAGTGGTATTCATGCTC | | |
| Synj1 | Synaptojanin 1 | CTGGAAGTTTGCGTGTTTCT | GTCTGTGAGCGTCTGGTTCT | | |
| Synpo2 | synaptopodin 2 | GGACAAGATGGAAATGTTGC | GTCCCATCTGGAAGGAACTT | | |
| Syp | synaptophysin | GGAGTGGGTCGATGTGACTT | TTGGCTGACTGGTCCTCTCT | | |
| Syt1 | synaptotagmin I | ACTCCAACCCCTACGTCAAG | CTTTCCCAATCACACAGTGG | | |
| Th | tyrosine hydroxylase | CTTCAGAAGAGCCGTCTCAG | GAATACCACAGCCTCCAATG | | |
| Tnf | tumor necrosis factor | CCCACTCTGACCCCTTTACT | TTTGAGTCCTTGATGGTGGT | | |
| Tnfrsf21 | tumor necrosis factor receptor superfamily, member 21 | CCACCACAGACACATTCTGA | TGGAGCTCTTTCGGATACTG | | |
| Tnfsf10 | tumor necrosis factor (ligand) superfamily, member 10 | ATTGGGAAGTCAGACCTGGA | TCATCCGTCTTTGAGAAGCA | | |
| Trp53 | transformation related protein 53 | AGCTTTGAGGTTCGTGTTTG | GGAACATCTCGAAGCGTTTA | | |
| Tubb4 | tubulin beta 4A class IVa | CAGCCTCCAAGTCCTCTCAC | TTCCTCGTCCTTCAGCACTT | DOWN | YES |
| Vamp1 | vesicle-associated membrane protein 1 | AATGTCTGCTCCAGCTCAAC | CCAAGACCTTGTCCACATTC | | |
| Vamp2 | vesicle-associated membrane protein 2 | TGAGGTTCCCATCACCTCTC | CTGTGGGGTTTGCTTTTGTT | | |
| Xiap | X-linked inhibitor of apoptosis | TGTGGACATCTGGTCACTTG | CTTAAAGTTCGCTCCCATCA | DOWN | NO |

### Slide 4
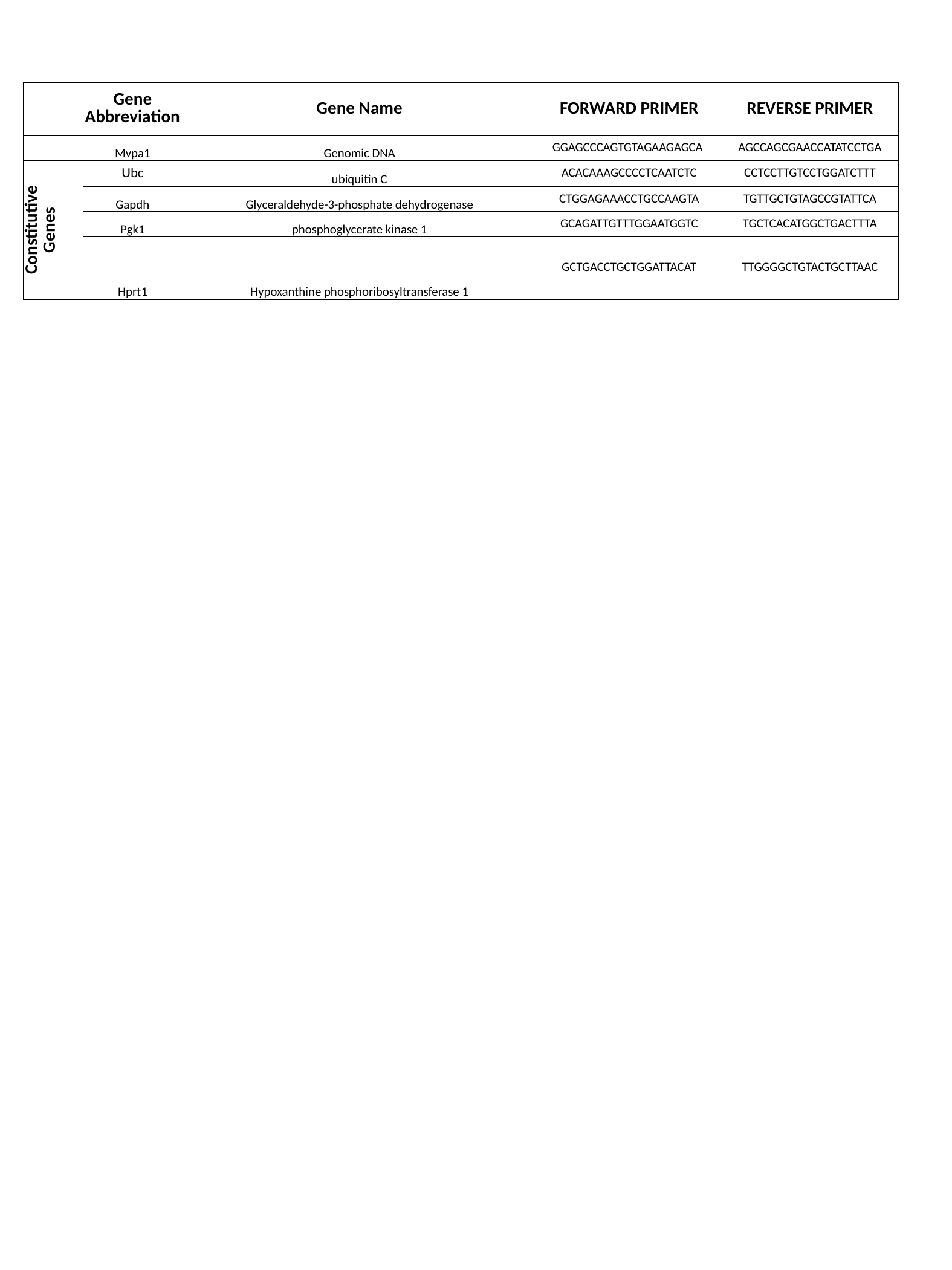

| | Gene Abbreviation | Gene Name | FORWARD PRIMER | REVERSE PRIMER |
| --- | --- | --- | --- | --- |
| | Mvpa1 | Genomic DNA | GGAGCCCAGTGTAGAAGAGCA | AGCCAGCGAACCATATCCTGA |
| Constitutive Genes | Ubc | ubiquitin C | ACACAAAGCCCCTCAATCTC | CCTCCTTGTCCTGGATCTTT |
| | Gapdh | Glyceraldehyde-3-phosphate dehydrogenase | CTGGAGAAACCTGCCAAGTA | TGTTGCTGTAGCCGTATTCA |
| | Pgk1 | phosphoglycerate kinase 1 | GCAGATTGTTTGGAATGGTC | TGCTCACATGGCTGACTTTA |
| | Hprt1 | Hypoxanthine phosphoribosyltransferase 1 | GCTGACCTGCTGGATTACAT | TTGGGGCTGTACTGCTTAAC |

### Slide 5
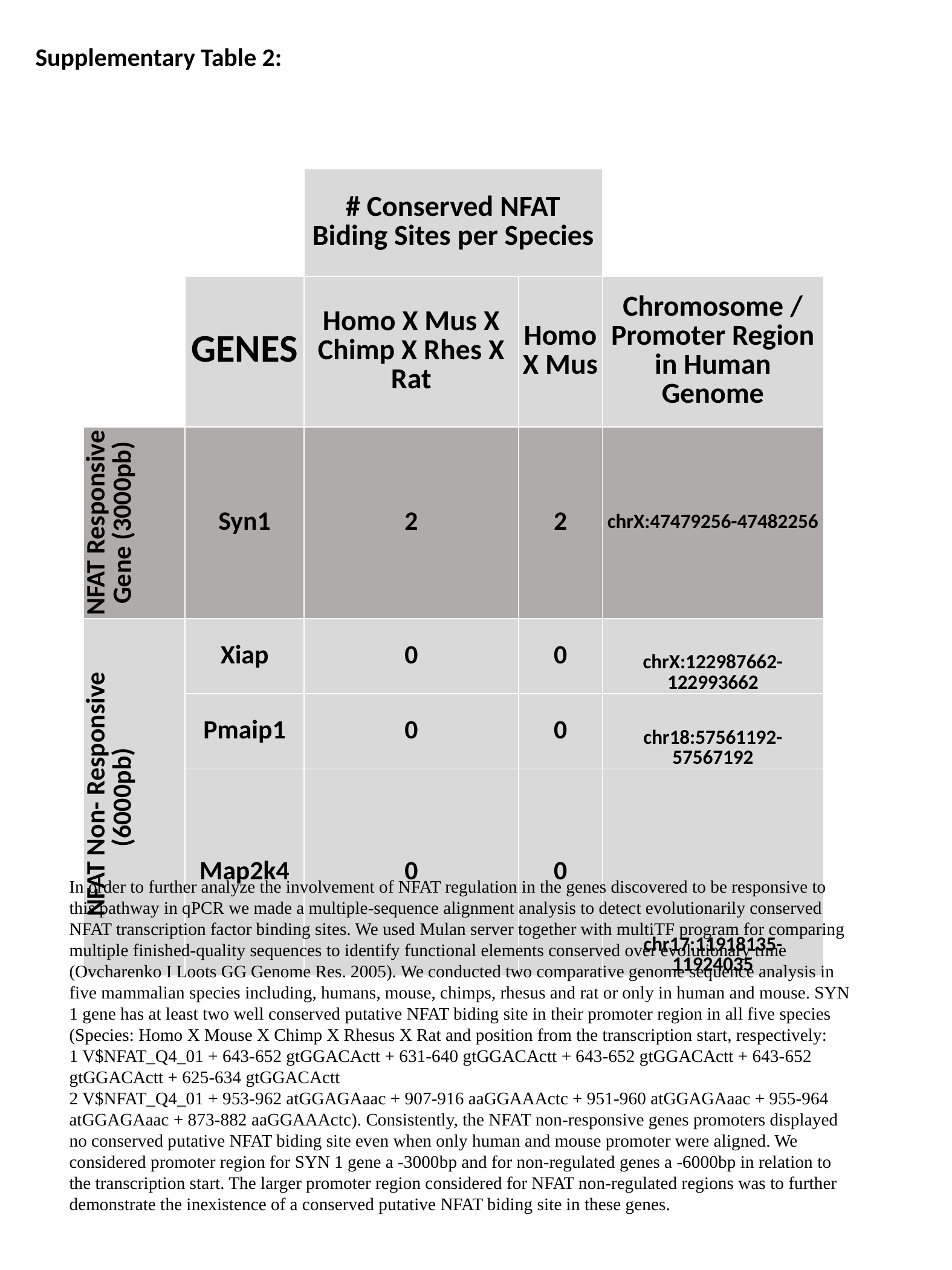

Supplementary Table 2:
| | | # Conserved NFAT Biding Sites per Species | | |
| --- | --- | --- | --- | --- |
| | GENES | Homo X Mus X Chimp X Rhes X Rat | Homo X Mus | Chromosome / Promoter Region in Human Genome |
| NFAT Responsive Gene (3000pb) | Syn1 | 2 | 2 | chrX:47479256-47482256 |
| NFAT Non- Responsive (6000pb) | Xiap | 0 | 0 | chrX:122987662-122993662 |
| | Pmaip1 | 0 | 0 | chr18:57561192-57567192 |
| | Map2k4 | 0 | 0 | chr17:11918135-11924035 |
In order to further analyze the involvement of NFAT regulation in the genes discovered to be responsive to this pathway in qPCR we made a multiple-sequence alignment analysis to detect evolutionarily conserved NFAT transcription factor binding sites. We used Mulan server together with multiTF program for comparing multiple finished-quality sequences to identify functional elements conserved over evolutionary time (Ovcharenko I Loots GG Genome Res. 2005). We conducted two comparative genome sequence analysis in five mammalian species including, humans, mouse, chimps, rhesus and rat or only in human and mouse. SYN 1 gene has at least two well conserved putative NFAT biding site in their promoter region in all five species (Species: Homo X Mouse X Chimp X Rhesus X Rat and position from the transcription start, respectively:
1 V$NFAT_Q4_01 + 643-652 gtGGACActt + 631-640 gtGGACActt + 643-652 gtGGACActt + 643-652 gtGGACActt + 625-634 gtGGACActt
2 V$NFAT_Q4_01 + 953-962 atGGAGAaac + 907-916 aaGGAAActc + 951-960 atGGAGAaac + 955-964 atGGAGAaac + 873-882 aaGGAAActc). Consistently, the NFAT non-responsive genes promoters displayed no conserved putative NFAT biding site even when only human and mouse promoter were aligned. We considered promoter region for SYN 1 gene a -3000bp and for non-regulated genes a -6000bp in relation to the transcription start. The larger promoter region considered for NFAT non-regulated regions was to further demonstrate the inexistence of a conserved putative NFAT biding site in these genes.
